# A lightweight, ultrafast and general embedding framework for large-scale spatial omics data

**DOI:** 10.64898/2026.02.04.703814

**Authors:** Bingjie Dai, Yuchao Liang, Litai Yi, Pengwei Hu, Yancheng Song, Binzhi Qian, Miaoxia He, Linhui Wang, Zhiyuan Yuan, Yongchun Zuo

## Abstract

Deciphering ultra-large-scale omics data with minimal resources while maintaining high computational efficiency is a longstanding challenge in biology. Here, we present Local Pooling (LP), a lightweight, ultrafast and general framework that leverage neighbor-indexing strategy and local pooling module to generate omics embedding and compatible with variety of downstream analyses. We developed its adaptation for spatial omics, SpaLP, and evaluated it on over 20 large-scale datasets spanning 9 technology platforms. SpaLP consistently outperformed baseline methods across multiple benchmarks, including niche identification, expression reconstruction, multiple slices integration, 3D organ construction, multi-omics integration and cross-platform generalization. Notably, SpaLP processed a 1.35-million-cell mouse embryo slice in just 47 seconds, achieving up to 300-fold increase in computational efficiency compared to Graph Neural Network (GNN)-based methods. Meanwhile, SpaLP increased the average adjusted rand index (ARI) by over 30% for niche identification in simulated and realistic settings. Furthermore, we applied SpaLP to integrate 8.4 million mouse brain cells within 4 minutes on a single GPU and constructed a 3D spatial atlas. Finally, we explored SpaLP’s ability of cross-platform generalization and potential for developing an omics foundation model. As a novel and general framework, we believe that LP could help more researchers develop new model on large-scale data and overcome the research barriers caused by computing resources in more fields.

## Introduction

Advances in high resolution spatial omics have facilitated the exploration of intricate heterogeneity at single-cell level. Compared to low-resolution technologies^1-6^, high-resolution omics technologies provide a more accurate representation of the cellular composition of the microenvironment, such as MERFISH^7^, CosMx^8^, MERSCOPE, Xenium^9^, Visium HD^10^, Stereo-seq^11^, Stereo CITE-seq^12^, STARmap PLUS^13^ and CODEX^14^. However, the rapid development of technology has significantly increased the scale and sparsity of data. Millions to tens of millions of cells pose new computational challenges for various tasks. Furthermore, there are significant differences in resolution and sequencing depth between the technical platforms, and tissue themselves also introduce different spatial distributions and batch effects. Numerous of computational approaches^15-24^ have been proposed to model spatial information for analysis of spatial omics data. Among them, graph-based framework, particularly graph neural networks (GNNs) methods have become the mainstream^25^. However, these methods are limited by GNN architectures that cannot handle large-scale tasks^26^. Despite a few methods like NicheCompass^27^ and HERGAST^28^ employ sampling strategies or Divide-Iterate-Conquer (DIC) approaches to address this issue, they still suffer from substantial computational resource and long runtimes. More importantly, these strategies will divide the complete tissue slice and disrupt the global graph structure, making it difficult to preserve the continuous spatial information essential for biological interpretation.

Here, we present Local Pooling (LP), a novel and general framework that efficiently generate omics embedding with extremely low computational resources and compatible with variety of downstream analyses. Unlike the adjacency matrix used in GNNs, LP employs an index matrix, allowing GPU memory usage to grow linearly with the number of graph nodes and significantly reducing memory consumption. This enables LP to handle data from tens of millions of cells only on a single GPU. Moreover, index matrix supports full graph input, avoiding the sampling destruction of the global structure in ultra-large-scale data. Based on the input of index matrix, LP efficiently aggregates neighborhood information via local pooling module. On million-scale datasets of spatial omics, LP achieves up to a 300-fold improvement in computational efficiency compared with GNNs. In addition, LP framework exhibits strong generalization capabilities. Only need a small amount of pre-training data, frozen parameters LP model can be directly applied to new data for inference. Given the generality of the adjacency matrix and index matrix, the LP framework can naturally be extended to more other fields.

For spatial omics, we developed SpaLP and evaluated its performance for various tasks on over 20 ultra-large-scale datasets spanning 9 technology platforms. We first demonstrated SpaLP’s performance in processing efficiency, niche identification and omics data reconstruction on simulated and experimentally acquired multiple ultra-large-scale samples, including the human lung cell atlas, human breast cancer, mouse brain, mouse spleen, mouse embryo and mouse testis. Notably, SpaLP processed a 1.35-million-cell mouse embryo slice in just 47 seconds, achieving up to 300-fold increase in computational efficiency compared to GNN-based methods. Meanwhile, SpaLP increased the average adjusted rand index (ARI) by over 30% for niche identification in simulated and realistic settings. We then evaluated SpaLP with competing methods for multi slices integration from same platform (MERSCOPE coronal mouse brain, STARmap PLUS sagittal mouse brain, Visium HD human tonsil and CODEX human tonsil) as well as across different platforms (CosMx, MERFISH and STARmap PLUS mouse brain; CosMx, Xenium and Visium HD Colorectal cancer). We also integrated 8.4 million mouse brain cells within 4 minutes on a single GPU and constructed a 3D spatial atlas to demonstrated its extremely low resource requirements. Furthermore, SpaLP successfully decoded the tumor microenvironment on a low-quality kidney cancer sample and captured more fine-grained histological details on a gastric cancer sample. SpaLP also achieved large-scale spatial multi-omics integration compared to GNN-based methods like SpatialGlue^21^ and COSMOS^23^. Finally, we explored SpaLP’s ability of cross-platform generalization and potential for developing an omics foundation model.

## Result

### The architecture of LP

SpaLP processes cell-level or bin-level spatial omics data by constructing a spatial index matrix that represents local cell neighborhoods, where each row corresponds to a central cell or bin and each column to its neighboring cells (Fig. 1a). Guided by the precomputed neighbor indices, each iteration of the LP architecture operates only within local neighborhoods, the cumulative effect of multiple iterations enables information to expand outward and cover the entire graph (Extended Fig. 1a). This capability enables SpaLP to generate embeddings for ultra-large-scale spatial omics slice with high computational efficiency. By concatenating index matrices derived from the neighbor graphs of different slices, SpaLP can further integrate features across slices into a shared latent space (Fig. 1a). We evaluated these advantages using datasets that cover a range of omics modalities and sequencing platforms, illustrating the broad applicability of LP in spatial omics.

**Fig. 1.**
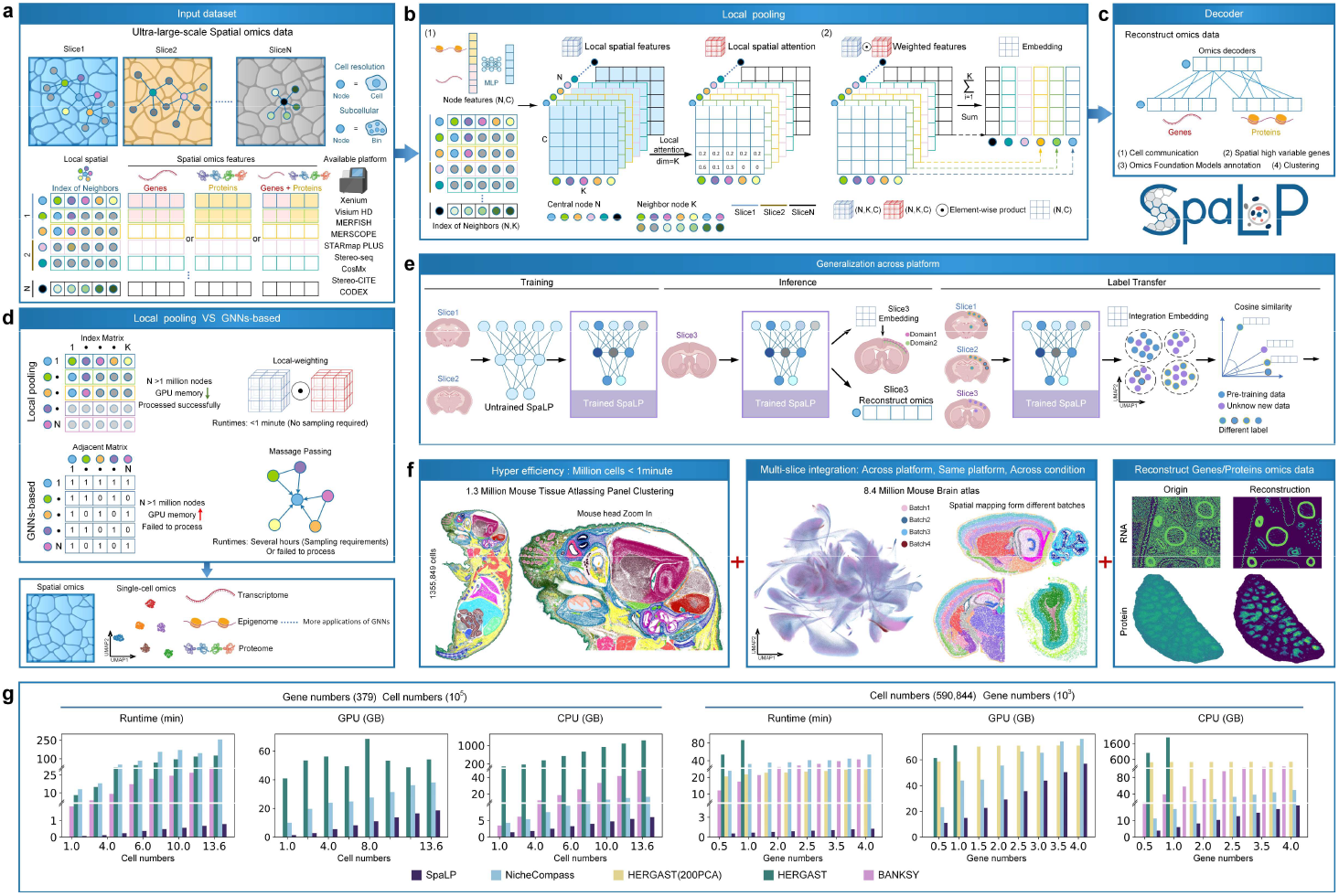
Overview of SpaLP. **a**, SpaLP accepts single slice or multi slices ultra-large-scale spatial omics data with cell or subcellar resolution observations as input. The spatial neighborhood graph was constructed by two-dimensional coordinates and converted into an index matrix, where the index represents each central cell and the corresponding matrix value is the neighbor index of the cell. SpaLP supports data from different omics platforms as input. **b**, Local pooling module can be divided into two phases: (1) The first stage uses MLP to individually extract its molecular signature for each cell. The node features with shape (N, C) and the index matrix with shape (N, K) are constructed as a 3D local spatial feature with shape (N, K, C). The neighborhood attention of local spatial features is calculated to obtain local spatial attention. (2) In the second stage, the weighted features are obtained by multiplying local spatial attention and features element-by-element, and then the weighted features of the neighborhood dimensions are accumulated and summed to obtain the microenvironment features of the central node. **c**, Decoders reconstruct omics counts and aggregated counts of its neighborhood. **d**, Compared with the GNN-based methods, the local pooling significantly reduces the GPU memory requirements, greatly improves the computational efficiency, and eliminates of the sampling requirements in ultra-large-scale data. Given the generality of adjacency matrix and index matrix, local pooling can be naturally extended to single cells and other fields where GNN can be applied. **e**, SpaLP has cross-platform generalization ability and can complete zero-shot inference. **f**, SpaLP facilitates downstream applications in spatial omics data analysis. **g**, Using different cell and gene scales to teste runtime, GPU and CPU memory of SpaLP and baseline methods.

To encode features of local neighbors, LP begins by applying a multilayer perceptron (MLP) to extract the individual information for each node. After obtaining embeddings of all nodes, index matrix is used to construct a local spatial feature (Fig. 1b), which represents a group of niche features (Extended Fig. 1a). For each feature dimension, we applied a local attention mechanism to assign attention weights to each neighbor node. The embedding of each node is obtained by summing the weighted feature matrix of the local neighborhood corresponding to each central node, and is decoded to reconstruct omics information (Fig. 1c).

The scale of index matrix grows linearly with the number of graph nodes, whereas the adjacency matrix expands quadratically in GNN. This enables LP to significantly reduce the resource consumption, which is pronounced when node population exceeding millions (Fig. 1d). Notably, iterations of local-weighting perform a function analogous to message passing, effectively aggregating information from neighboring nodes. Compared with GNNs, LP achieves a computation speed hundreds of times faster on large-scale datasets, while enabling full-graph input. Given the generality of the adjacency matrix and index matrix, the LP framework can naturally be extended to more other fields.

SpaLP compatibles with variety of downstream analyses (Fig. 1e, f), including niche identification, expression reconstruction, multiple slices integration, 3D organ construction, multi-omics integration and cross-platform generalization. We demonstrated the superior computational efficiency and resource utilization of SpaLP across datasets with varying cell numbers and gene counts (Fig. 1g). For instance, SpaLP processes 1.35 million cells in just 47s, while the second-fastest method, BANKSY^29^, takes 30 minutes. In comparison, both the GNN-based method NicheCompass and HERGAST require significantly runtime and resources. NicheCompass takes over four hours, while HERGAST is unable to handle million-scale data due to memory requirements exceeding 1,200 GB. We further comprehensively evaluated SpaLP on diverse tissues and species, covering tasks comprising tens of thousands to nearly ten million cells (Extended Data Fig. 1b).

### Benchmarking SpaLP and existing methods

We benchmarked the performance of SpaLP on large-scale spatial omics niche identification using both simulated and experimentally acquired data. For datasets with ground-truth annotations, we assessed model performance using six supervised metrics. For experimentally acquired datasets without annotations, we compared the performance of SpaLP and baseline methods based on their ability to identify specific regions. As our primary focus is on evaluating performance at scales exceeding 100,000 cells, we compared SpaLP against three state-of-the-art methods specifically designed for large-scale spatial omics: NicheCompass, HERGAST, and BANKSY.

We first evaluated it on a simulated dataset with 640,000 cells generated by Gong et al.^28^, which features distinct spatial patterns and complex, fine-grained spatial structures, together with ground-truth annotations (Fig. 2a). SpaLP efficiently processed the simulated dataset in just 49s, whereas BANKSY required approximately 29 minutes, HERGAST took 31 minutes, and NicheCompass needed 1 hour and 45 minutes (Fig. 2b). In addition to its superior efficiency, SpaLP outperformed all other methods in accuracy (ARI = 0.9), and precisely captured the structural organization of each spatial region, while baseline methods exhibited varying degrees of discrete distribution across several regions (Fig. 2c and Supplementary Fig. 1). SpaLP was also capable of reconstructing gene expressions with significant spatial patterns (Fig. 2d, Supplementary Fig. 2, and Extended Data Fig. 2a, b). To evaluate the reconstruction performance of SpaLP, we employed the omics foundation model SCimilarity^30^ to generate embeddings for both the original and reconstructed gene expression profiles. The result showed the reconstructed expression more accurately reflected the cell type differences (Extended Data Fig. 2c). We performed a grid search over multiple leiden resolution settings to further evaluate the robustness of SpaLP. Across different niche number partitions, SpaLP consistently maintained stable and superior performance (Fig. 2e).

**Fig. 2.**
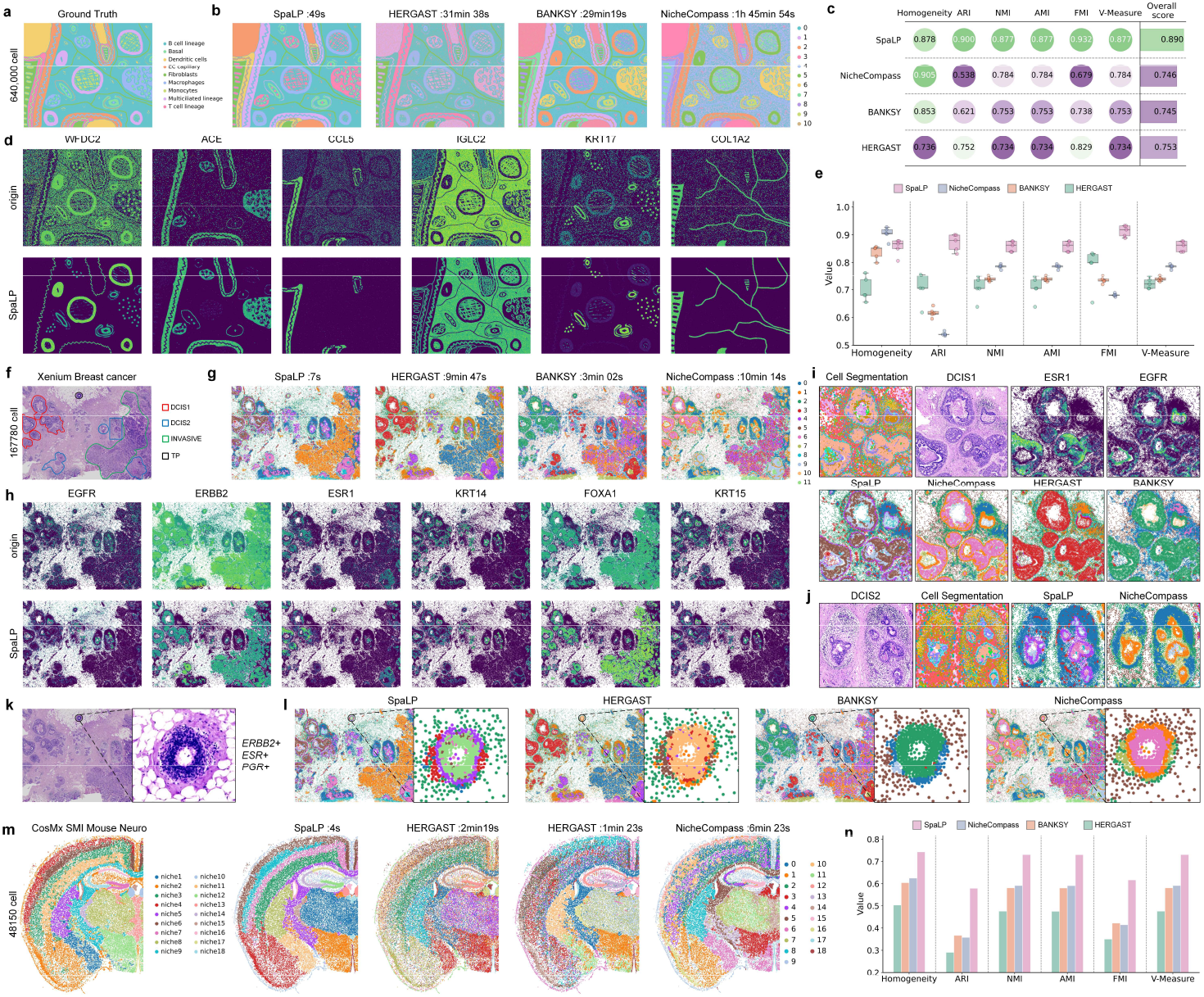
The performance of SpaLP in single-sample niche identification. **a**, Ground Truth of simulation data. **b**, Visualization of the niche identification results for SpaLP and baseline methods on the simulation data. **c**, Quantitative evaluation of four methods with six supervised metrics. **d**, Spatial mapping of original and reconstructed genes expression. **e**, Box plots of six supervised metrics with scores of clustering results with the number of clusters ranging from 9 to 13. In the box plot, the center line denotes the median, box limits denote the upper and lower quartiles, and whiskers denote the 1.5 times the interquartile range. **f**, H&E image and annotation of pathological regions from original study. **g**, Visualization of the niche identification results for SpaLP and baseline methods on the breast cancer data. **h**, Spatial mapping of original and reconstructed genes expression. **i**, Visualization of the DCIS1 region cell segmentation from 10X Genomics reports and niche identification results from four methods. **j**, Visualization of the DCIS2 region cell segmentation from 10X Genomics reports and niche identification results from two methods. **k**, Visualization of the triple-positive (TP) region of the HE images magnification results. **l**, Visualization of the niche identification results for SpaLP and baseline methods on the triple-positive (TP) region. **m**, Spatial regions of ground truth provided by NanoString CosMx. **n**, Visualization of the niche identification results for SpaLP and baseline methods on the mouse brain data. **o**, Quantitative evaluation of four methods with six supervised metrics.

To investigate the SpaLP’s ability to capture complex microenvironments of tumor tissue^31^, we applied it to a Xenium breast cancer slice^9^ (167,780 cells, 313 genes). This dataset has been annotated to distinguish between invasive cancer, ductal carcinoma in situ DCIS1 and DCIS2 (Fig. 2f). Consistent with previous observations, SpaLP processed the entire dataset in just 7s (Fig. 2g) and the reconstructed expression demonstrated a stronger spatial pattern (Fig. 2h). SpaLP also showed the highest consistency with the cell segmentation results of 10x Genomics reports in two DCIS regions (Fig. 2i, j). Notably, in the HERGAST signal amplification test, ESR1 and EGFR were reported as two genes with complementary spatial distributions (Extended Data Fig. 2d). This distribution was highly overlap with the two independent niches identified by SpaLP. Based on previous studies^9^ and the expression of breast cancer genes (Extended Data Fig. 2e), we identified the two niches as Estrogen Receptor-positive region (ER+) and Triple-negative region (TN) (Extended Data Fig. 2f). Furthermore, SpaLP also recognized a special niche that was not separately delineated in the results of BANKSY and NicheCompass (Fig. 2k, l). This niche had been designated as the Triple-positive region (TP) in prior studies^9^ (Extended Data Fig. 2g), and spatial gene expression mapping validated this designation (Extended Data Fig. 2e). These findings proved SpaLP can accurately resolve complex tumor microenvironments, as it is the only method capable of identifying all three distinct regions: ER^+^, TN, and TP.

To further evaluate SpaLP across multiple technologies and tissue contexts, we compared it with baseline methods on a NanoString CosMx mouse brain dataset (48,150 cells, 950 genes) and three CODEX mouse spleen datasets (∼80,000 cells, 30 proteins). The results showed that SpaLP accurately identified cortical subdivisions (Fig. 2m) and outperformed baseline methods across six metrics (Fig. 2n). The reconstructed gene also exhibited significant spatial patterns (Extended Data Fig. 2h, i). In the CODEX proteomic datasets, SpaLP consistently demonstrated leading performance, achieving the best niche identification across all three mouse spleen datasets (Extended Data Fig. 3).

### Precisely deciphering anatomy across different tissues

We further applied SpaLP to datasets from mouse embryo, testis, and spleen to display its resolving power across diverse tissue types. On a Xenium whole mouse pup slice (∼1.35 million cells), SpaLP completed the processing within 47s, requiring only 16 GB of GPU memory and 6 GB of CPU memory. BANKSY required approximately 31 minutes, whereas NicheCompass took more than 4 hours. Although HERGAST effectively reduces GPU memory usage through its DIC strategy, it still requires over 1,200 GB of CPU memory to run this dataset, which makes it impractical for million-scale data processing (Fig. 3a and Supplementary Table S1).

**Fig. 3.**
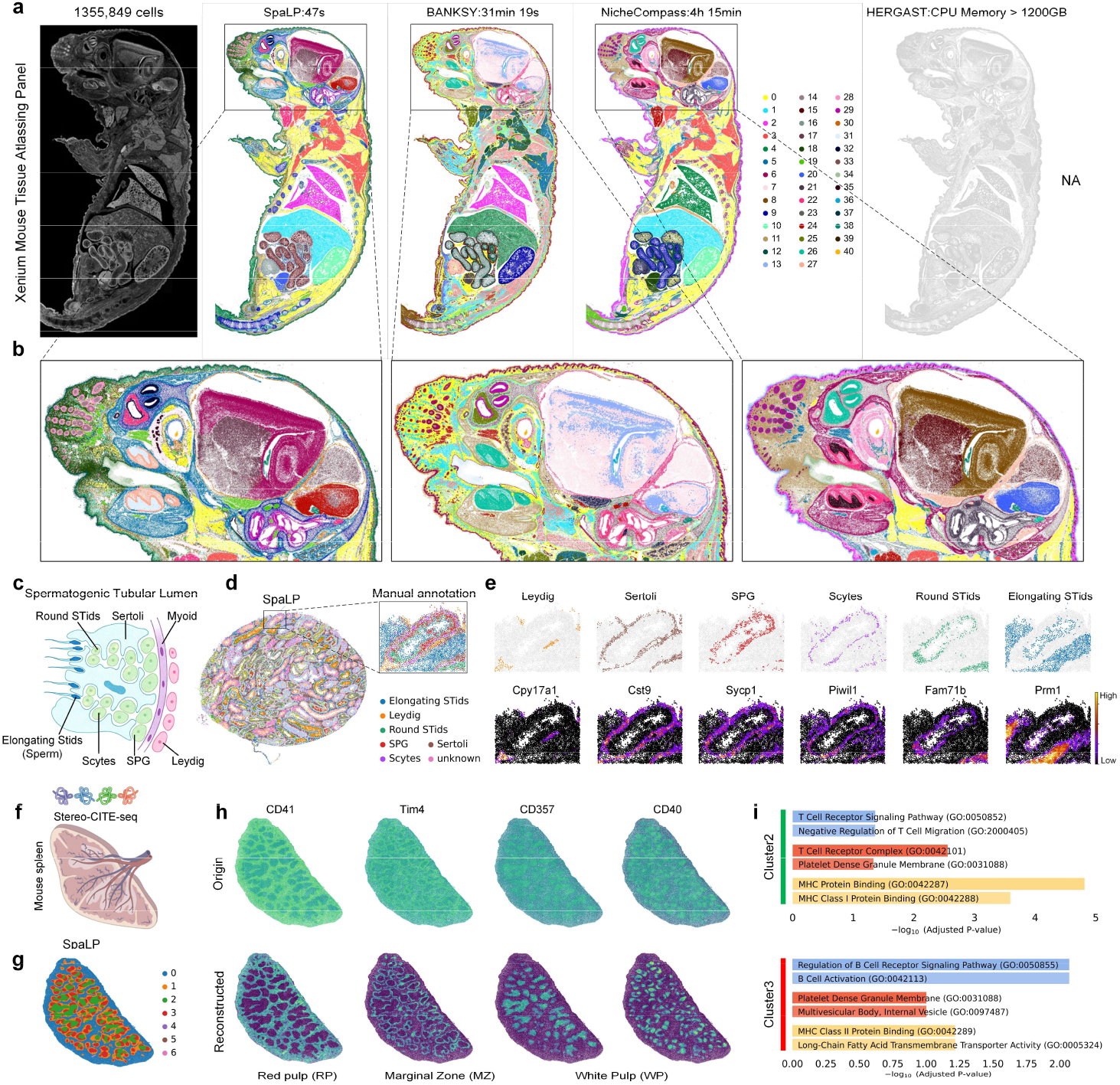
Deciphering anatomy across different mouse tissues and modalities using SpaLP. **a**, Visualization of the niche identification results for SpaLP and baseline methods on whole mouse pup. The final result was not available because HERGAST required more than 1200GB of CPU Memory. **b**, Visualization of the niche identification results for three methods on the mouse head. **c**, Distribution of cell types in the Spermatogenic Tubular Lumen. **d**, Niche identification results for SpaLP with a zoomed-in view of a single seminiferous tubule region. **e**, Highlighted visualization of the ring-like niches and the corresponding marker gene expression. **f**, SpaLP resolves mouse spleen using single cell spatial proteome. **g**, Visualization of the niche identification results for three methods on the mouse spleen. **h**, Spatial mapping of original and reconstructed expression for selected marker genes from different anatomical regions. **i**, Gene Ontology analyses in niches 2 and 3 identified by SpaLP on Stereo CITE-seq mouse spleen.

We then selected two regions of interest (ROIs) in the mouse head to evaluate the performance of SpaLP (Fig. 3b and Extended Data Fig. 4a). Among the results of the three algorithms, only SpaLP distinguished precise niches within the two ROIs, whereas the corresponding regions in the outputs of BANKSY and NicheCompass exhibited a uniformly smoothed result (Extended Data Fig. 4b). We visualized the expression of the marker genes *Slc22a6* and *Runx2* in this region and found that their spatial distributions were highly consistent with the results identified by SpaLP (Extended Data Fig. 4c). Similar patterns were observed across multiple regions of the mouse brain, where the spatial expression of the corresponding marker gene was consistent with niches identified by SpaLP (Supplementary Fig. 3). These results showed that SpaLP provided more precise decipherment for large-scale slices.

To examine the ability of SpaLP to capture special spatial patterning at high resolution, we utilized a mouse testis dataset generated using the Stereo-seq platform (198,248 cells, 27,869 genes). In mammals, germ cells at successive developmental stages are arranged in a radial or ring-like gradient toward the lumen, surrounded by supporting Sertoli cells and interstitial Leydig cells (Fig. 3c). A complete mouse testis slice can reveal the clear developmental progression during spermatogenesis^32-36^. SpaLP identified the densely packed tubular architecture as well as the ring-shaped, lumen-oriented gradient niches that represent the seminiferous tubules (Fig. 3d). We then zoomed in on a complete seminiferous tubule region and annotated ring-like niches using markers from previous studies^37^ (Fig. 3e and Extended Data Fig. 4d, e). SpaLP accurately resolved the circular arrangement of male germ cells, where the major cell populations essential to spermatogenesis were positioned along the same tubule in an order consistent with known anatomical patterns, including Leydig cells, Sertoli cells, SPG, spermatocytes (Scytes), round spermatids (Round STids), and elongating spermatids^38-43^ (Elongating Stids). In contrast, traditional embedding method failed to capture hierarchical structures (Extended Data Fig. 4f, g). Meanwhile, visualization of reconstructed gene distributions revealed highly specific spatial patterns consistent with the observable regional arrangement (Extended Data Fig. 4e and Supplementary Fig. 4). These findings support the ability of SpaLP to analyze slices with complex hierarchical structures.

We next applied SpaLP to a mouse spleen dataset from Stereo CITE-seq (295,215 cells, 128 proteins), for demonstrating its robustness on spatial proteomics data with substantially lower feature dimensionality^44,45^ (Fig. 3f). The densely packed tissue architecture and complex arrangement of immune populations in the spleen pose challenges to identifying spatially coherent niches^46-48^. SpaLP accurately delineated spatial niches highly concordant with established splenic anatomy^49^ (Fig. 3g and Supplementary Fig. 5a). Moreover, expression distributions reconstructed by SpaLP exhibited high specificity across distinct niches (Fig. 3h). In the white pulp, SpaLP further resolved two distinct subniches and we defined them as T cell-enriched and B cell-enriched regions based on their corresponding markers and previous studies^1,50-52^ (Supplementary Fig. 5c, d). Gene Ontology (GO) analysis further confirmed this conclusion (Fig. 3i). The reconstructed expression of multiple markers also revealed strong concordance with previously observed patterns (Supplementary Fig. 5d-f).

### Integrating multiple ultra-large-scale slices across platforms

We systematically evaluated the capability of SpaLP to integrate multiple slices across diverse scenarios, including within and cross platform integration. For the within-platform integration benchmark, we first applied SpaLP to a MERSCOPE mouse brain atlas dataset comprising 9 slices, 734,686 cells, and 483 genes. SpaLP significantly outperformed existing methods in computational efficiency (Fig. 4a). Two spatial consistency metrics, Cell-type Affinity Similarity (CAS) and Graph Connectivity Similarity (GCS), revealed that the full-graph input of the LP framework better preserved spatial structures than baseline methods. Multiple niches were identified across 9 slices using annotations from three coronal planes in the Allen Brain Atlas: S1, S2, S3 (Fig. 4b, c and Supplementary Fig. 6a). SpaLP outperformed both NicheCompass and BANKSY in achieving the most consistent and spatially coherent niche delineation across multiple brain regions, including the Thalamus, Caudoputamen, Corpus Callosum, Isocortex Layers, Hippocampal region and Pontine Gray (Supplementary Fig. 6b). We subsequently applied it to three sagittal slices from the STARmap PLUS platform (3 slices, 422,673 cells, 1022 genes), the result showed SpaLP efficiently revealed consistent and spatially coherent niches (Fig. 4d, e).

**Fig. 4.**
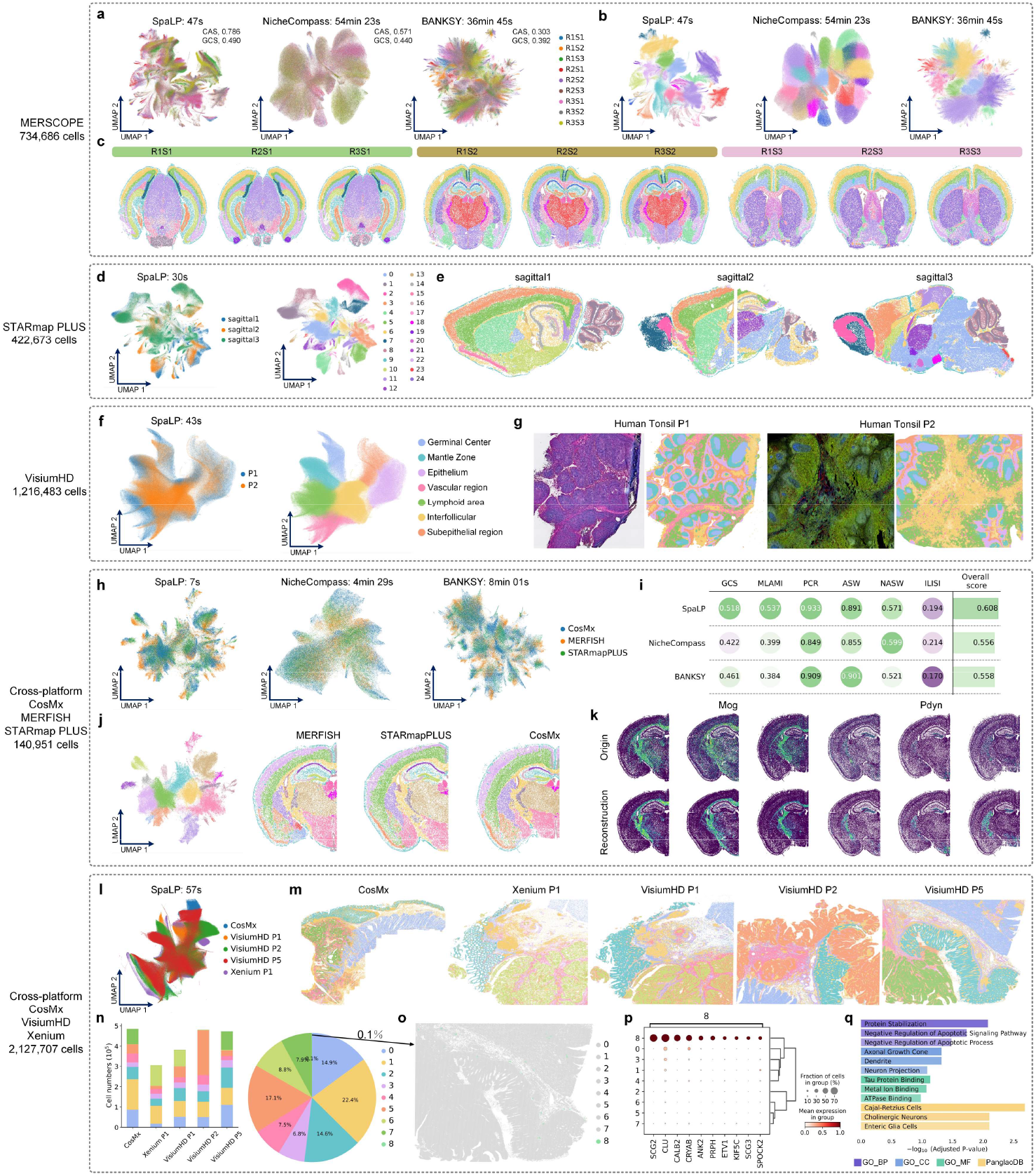
The performance of SpaLP in multi-slices integration. **a**, Integrated embedding Uniform manifold approximation and projection (UMAP) from SpaLP and baseline methods colored by 9 coronal mouse brain slices source. **b**, UMAPs colored by niche identification results from SpaLP and baseline methods. **c**, Visualization of the niche identification results for SpaLP on 9 coronal mouse brain slices. **d**, SpaLP’s integrated embedding UMAP and colored by 3 sagittal slices and the niche identification results. **e**, Visualization of the niche identification results for SpaLP on 3 sagittal mouse brain slices. **f**, SpaLP’s integrated embedding UMAP and colored by 2 human tonsil slices and the niche identification results. **g**, H&E and Immune fluorescence images from 10X Genomics reports and the niche identification results for SpaLP on 2 human tonsil slices. **h**, SpaLP’s integrated embedding UMAP and colored by 3 platform mouse slices. **i**, Quantitative evaluation of three methods with six metrics. **j**, UMAPs and spatial mapping colored by niche identification results from SpaLP on three different platforms. **k**, Spatial mapping of original and reconstructed genes expression. **l**, SpaLP’s integrated embedding UMAP and colored by 5 colorectal cancer slices from 3 platforms. **m**, Visualization of the niche identification results for SpaLP on 5 colorectal cancer slices. **n**, Bar plot of niche distribution for each colorectal cancer slice, and pie plot of niche distribution for all slices. **o**, Highlighted visualization of the rare niche. **p**, Dot plot of the highly expressed genes in SpaLP’s unique niches. **q**, Gene Ontology and cell type enrichment analyses.

Next, we applied SpaLP to two Visium HD human tonsil slices (1,216,483 cells, 18,085 genes, per slice encompasses ∼500,000 cells) to recognize key niches in tissues lacking stereotypical spatial patterns. SpaLP integrated two ultra-large-scale slices within 43s and identified seven niches (Fig. 4f), which were delineated based on stained images and previous research^53^. As two key regions of the tonsil, the germinal center and mantle zone were accurately divided (Supplementary Fig. 7a, b). In addition, reconstructed gene expression exhibited more pronounced and spatially coherent patterns in multiple slices (Supplementary Fig. 7c). We then applied SpaLP to three CODEX tonsil slices (1,395,992 cells, 43 proteins) to identify the same regions in proteomics data. SpaLP maintained outstanding efficiency (Supplementary Fig. 8a), with its results showing high consistency with annotations from the original research^54^ (Supplementary Fig. 8b, c). The germinal center and mantle zone were also accurately identified in the B cells regions (Supplementary Fig. 8d). This finding demonstrates the robustness of SpaLP in integrating tissue slices and identifying essential niches across diverse omics data.

For the cross-platform integration, discrepancy in resolution, sequencing depth, and gene panels substantially increase the difficulties compared to within-platform integration. We firstly utilized three mouse brain slices from distinct platforms: STARmap PLUS (43,341 cells, 1,022 genes), MERFISH (49,430 cells, 1,122 genes), and CosMx (48,180 cells, 950 genes). SpaLP maintained highest computational efficiency and achieved the best performance in the comprehensive assessment across six metrics (Fig. 4h, i and Supplementary Note 1). Two global spatial conservation metrics (CAS and MLAMI) showed that the LP framework better preserved the spatial structure of the tissue, while the PCR score indicated that SpaLP effectively removed batch effects. Compared to baseline methods, SpaLP accurately captured mouse brain anatomy across different platforms and supported cross-platform reconstruction (Fig. 4j and Supplementary Fig. 9).

We then applied SpaLP to dissect complex microenvironments on 5 colorectal cancer (CRC) samples, involving the CosMx (493,834 cells, 18,000 genes), Xenium (307,762 cells, 422 genes), and Visium HD (three slices with 507,684, 545,913, and 541,968 cells, 18,085 genes). SpaLP integrated 2 million cells in 57s (Fig. 4l) and successfully identified conserved niches across different samples, such as the goblet cell-rich and fibroblast-rich niches (Extended Data Fig. 5a, b and Supplementary Fig. 10). Additionally, SpaLP preserved the variation in cancer regions across samples and classified them into three main types consistent with the original research^10^ (Extended Data Fig. 5c).

Notably, SpaLP identified a rare niche comprising only 0.1% of the 2 million cells (Fig. 4n). We then highlighted the spatial distribution of this niche in Visium HD P5 slice and extracted the highly expression genes (Fig. 4o, p). Neuron-related markers significantly enriched and aligned with the niche distribution (Extended Data Fig. 5d). Specifically, *SCG2, SCG3*, and *CRYAB* are involved in various functions of the enteric nervous system^55-58^. *PRPH* and *CALB2* have been reported as specific markers for enteric neurons^59-62^, and *KIF5C* encodes neuronal kinesin^63^. Meanwhile, GO and cell type enrichment analyses demonstrated that the gene set was significantly associated with neuronal cell type, structures, and functions (Fig. 4q). Based on these findings, we define the rare niche as an intestinal neuron enriched region. This experiment displayed that SpaLP accomplish effective integration on complex microenvironments, while preserving slice-specific biological variation and detecting rare niches.

### Constructing a whole-brain 3D atlas

We integrated a MERFISH whole-mouse-brain dataset comprising 8.4 million cells combined from 239 sections to demonstrated the low resource requirement of SpaLP. The process took only 3 mins and 40s on a single GPU with memory usage stayed around 24 GB. Moreover, SpaLP mapped all slices into a unified latent space and effectively eliminated batch effects (Fig. 5a). It accurately identified multiple niches that were consistent with the anatomical annotations of the Allen Brain Atlas (Fig. 5b, c). The major anatomical regions of the mouse brain were delineated into conserved niches that were spatially coherent (Fig. 5d, Extended Data Fig. 6a and Supplementary Fig.11-14). We visualized original and reconstructed expression of marker genes in the isocortex, hippocampus, cerebellum and striatum^64-71^. The result of SpaLP exhibited more significant spatial patterns and higher spatial autocorrelation scores, revealing the hierarchical structure across major regions (Fig. 5e, Extended Data Fig. 6b).

**Fig. 5.**
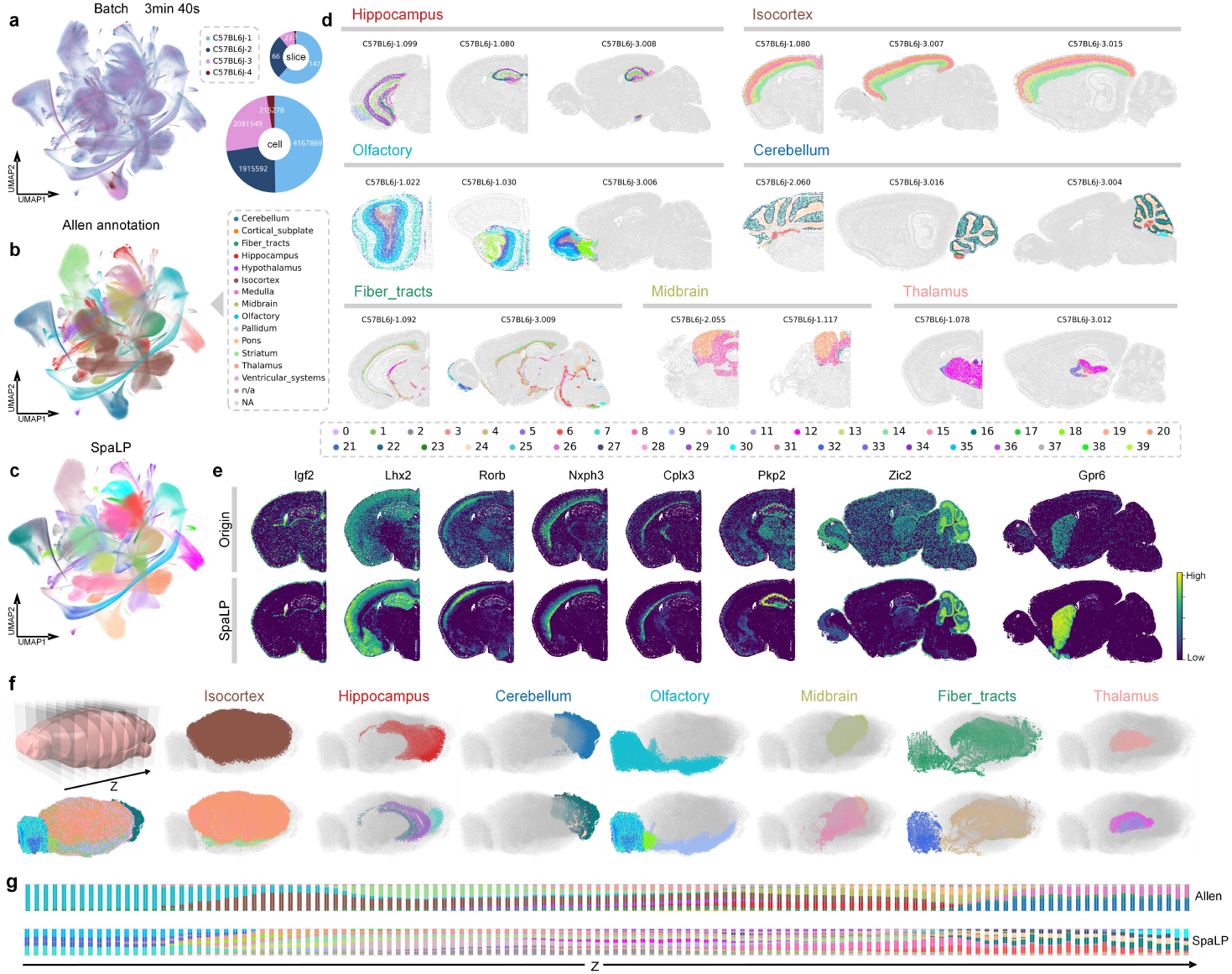
SpaLP integrates 8.4 million cells across 239 tissue slices and constructs a 3D atlas. **a**, SpaLP’s integrated embedding UMAP and colored by 4 batches. The right hollow ring plots show the distribution of the cell numbers and the distribution of the slice numbers in different batches. **b**, UMAP colored by annotation of mouse brain from the Allen Brain Atlas. **c**, UMAP colored by the niche identification results from SpaLP. **d**, Randomly selected tissue slices for each major brain region, colored by identified niches. **e**, Spatial mapping of original and reconstructed genes expression. **f**, 3D atlas generated from 129 consecutive coronal sections. Top 3D atlas colored by annotation of mouse brain from the Allen Brain Atlas. Bottom 3D atlas colored by identified niches. **g**, Niches distribution in 129 consecutive coronal sections. Top distribution colored by annotation of mouse brain from the Allen Brain Atlas. Bottom 3D atlas colored by identified niches.

At the whole-brain level, we constructed a 3D atlas of 129 consecutive coronal sections (from the olfactory bulb to the medulla oblongata), which consisted with the annotations of the Allen Brain Atlas (Fig. 5f and Extended Data Fig. 7). SpaLP revealed a continuous transition of niches across 3D space, displaying its potential for constructing 3D organ atlases (Fig. 5g and Supplementary Fig.15-22). In general, handling omics datasets at the tens-of-millions scale requires substantial computational resources and prohibitively long processing times^72,73^, making such analyses challenging for existing methods. The LP framework overcomes these limitations, showcasing strong promise for supporting the next generation of large-scale omics models.

### Robust niche identification across multiple cancer tissues

To evaluate the robustness of SpaLP across multiple human cancer tissue, we applied it to an in-house low-quality kidney cancer slice, an indistinct gastric cancer slice, and a multi-omics renal cell carcinoma (RCC) slice. For the in-house data, we performed NanoString CosMx Spatial Molecular Imager on a formalin-fixed, paraffin-embedded (FFPE) kidney cancer sample that had been preserved for three years, generating a dataset of 1,236,281 cells using a 1000-plex RNA panel across 1,037 fields of view (FOVs). Since RNA in long-term preserved FFPE samples degrades over time, it leads to reduced overall capture rates and heightened variability in capture rates across FOVs (Extended Data Fig. 8a).

Guided by hematoxylin-eosin (HE) staining and multiple immunofluorescence (mIF), we selected five regions to evaluate the performance of SpaLP and the baseline methods (Fig. 6a). SpaLP successfully captured fine-grained and coherent spatial niches, whereas NicheCompass and BANKSY generated either overly smoothed or grid-like structures (Fig. 6b). The niches identified by SpaLP in IRR, IAR, HSR, and InfSR regions exhibited high consistency with mIF staining (Fig. 6c, d) and the spatial mapping of marker genes (Extended Data Fig. 8b). We then annotated this slice using marker genes documented in prior studies^74,75^ (Extended Data Fig. 8c, d). The reconstructed expression of markers preserved identical spatial specificity patterns with the original data, while enhancing differences between niches. In the TER region, SpaLP precisely identified glomerulus and exhibited a more pronounced spatial distribution of gene at its core positions (Extended Data Fig. 8f).

**Fig. 6.**
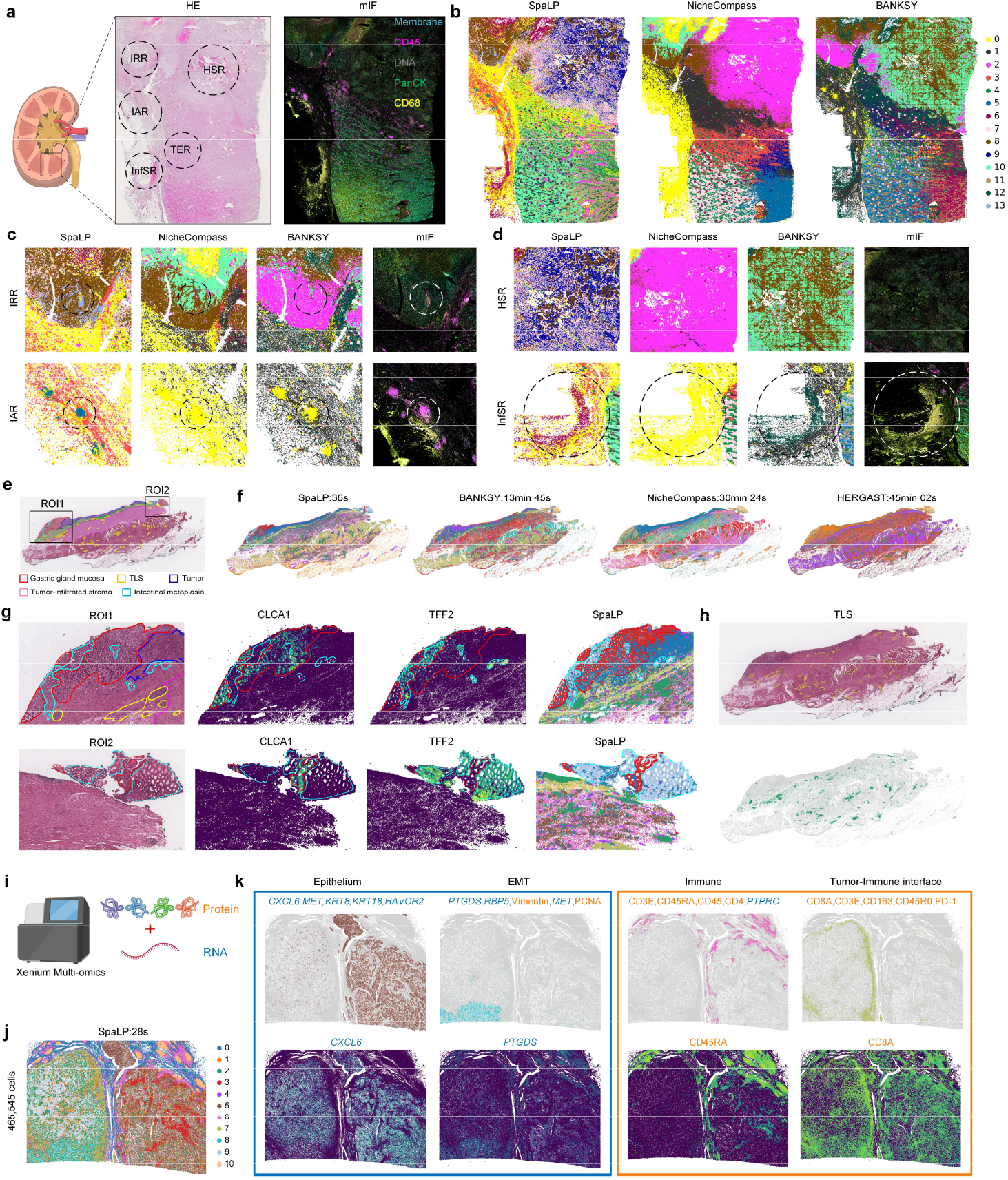
The performance of SpaLP in various cancer samples. **a**, H&E and multiple immune fluorescence (mIF) images. **b**, Visualization of the niche identification results for SpaLP and baseline methods on kidney cancer slice. **c**, Magnified visualization of three methods’ niches and mIF images in the IRR and IAR regions. **d**, Magnified visualization of three methods’ niches and mIF images in the HSR and InfSR regions. **e**, H&E image and annotation of pathological regions from original study. **f**, Visualization of the niche identification results for SpaLP and baseline methods on gastric cancer slice. **g**, Highlighted visualization of the Gastric gland mucosa and Intestinal metaplasia in two regions of interest (ROI1 and ROI2). CLCA1 and TFF2 are markers of the two annotation cell types. **h**, Highlighted visualization of the Tertiary Lymphoid Structure (TLS) and corresponding niche. **i**, Xenium Spatial multi-omics sequencing of transcriptomes and proteomes from the same slice. **j**, Visualization of the niche identification results for SpaLP on human renal cell carcinoma (RCC) slice. **k**, Visualization of four niches from SpaLP and corresponding gene or protein markers (Blue are genes and orange are proteins).

Next, we applied SpaLP to a Xenium gastric cancer data (696,314 cells, 377 genes) to evaluate its performance on indistinct microenvironment. Based on the previous research^76^, we delineated and annotated five distinct regions: Tumor, Tertiary Lymphoid Structure (TLS), Tumor-infiltrated stroma, Gastric gland mucosa and Intestinal metaplasia (Fig. 6e). SpaLP exhibited the most accurate identification of niches consistent with manual annotations (Fig. 6f). It successfully divided the niches of intestinal metaplasia and gastric gland mucosa in two ROIs (Fig. 6g). In contrast, BANKSY and HERGAST failed to identified the gastric gland mucosa, and NicheCompass incorrectly mixed them into a single niche (Supplementary Fig. 23). SpaLP also identified TLSs, which are key indicators of immune dynamics within the tumor microenvironment and are often associated with enhanced immune responses (Fig. 6h).

We used SpaLP on a large-scale Xenium renal cell carcinoma (RCC) tissue slice (465,545 cells, 396 genes, and 27 proteins), which is unable to process by traditional spatial multi-omics methods. The types of top markers in different niches displayed omics-specific preferences. Specifically, the top five markers in the epithelium were exclusively RNA, while those in the immune region were exclusively protein (Fig. 6i-k). These results demonstrated that SpaLP effectively leverages complementary information from different modalities to identify distinct niches within the multi-omics cancer tissue slice.

### Cross-platform generalization ability

Given the large-scale data integration capability of the SpaLP, we explored the potential of the LP framework to build an omics foundation model. We firstly pre-trained the SpaLP on tissue sections from multiple platforms, and then froze the model parameters (Fig. 7a). Next, we directly input new, unseen data into the trained model to inference embedding. We then transferred the label from pre-training data to each cell of new data with the highest similarity (Fig. 7b).

We initially utilized mouse coronal section datasets as pre-training data from three distinct platforms: CosMx, MERFISH, and STARmap PLUS (Fig. 7c). After pretraining and freezing the parameters, we input two datasets from MERFISH and Stereo-seq into trained SpaLP, respectively. The results indicated that the new data were effectively integrated with the pre-trained datasets (Supplementary Fig. 24a), and the transferred labels showed high consistency across different coronal sections. The cosine similarity analysis further validated the effective integration and provided interpretability for the prediction process (Fig. 7d).

**Fig. 7.**
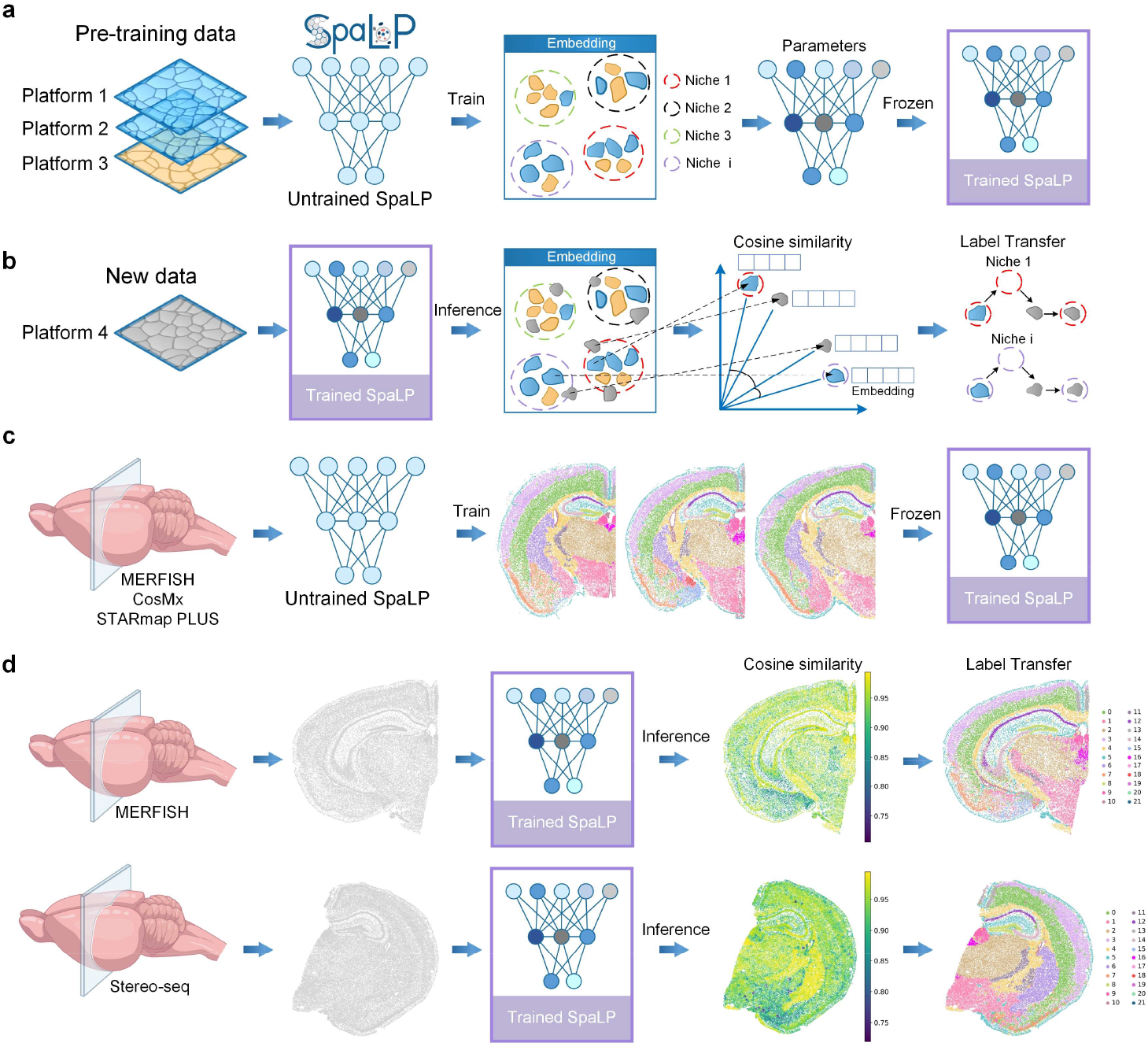
SpaLP can be generalized across platforms. **a**, SpaLP uses a small amount of data from different platforms for pre-training and freeze the trained model parameters. **b**, SpaLP is able to achieve zero-shot inference. Feeding unseen data into a trained SpaLP model to infer embedding and using cosine similarity to transfer the labels between new data and pre-training data. **c**, Models were pre-trained and niches identified using mouse coronal brain slices from MERFISH, CosMx and STARmap PLUS. **d**, New brain slices from the MERFISH and Stereo-seq platforms are input to the trained model to achieve label transfer. Cosine similarity scores provide an explanation for label transfer credibility.

In addition, we also expanded the experiment to the proteomics data. Specifically, we used three CODEX mouse spleen sections, two of which were used for pre-training and the other one as testing data. Consistent with the transcriptomic results, SpaLP effectively integrated the testing inference data and pre-trained data (Supplementary Fig. 24b), and achieved high-quality label transfer (Supplementary Fig. 25). These results showed the strong generalization ability of the LP framework, demonstrating the potential to develop omics foundation model.

## Discussion

We present LP, a lightweight, ultrafast, and general embedding framework to overcome the research barriers caused by limited computational resources in biology. Unlike previous methods, LP leverages the neighbor-indexing strategy and local pooling module to efficiently generate omics embedding with minimal resources. To validate the advantages of the LP framework, we applied its spatial omics adaptation, SpaLP, in comprehensive experiments. SpaLP processed a 1.35-million-cell mouse embryo slice in just 47s, and integrated 8.4 million mouse brain cells within 4 minutes on a single GPU using only 24 GB memory. Meanwhile, SpaLP increased the ARI by over 30% for niche identification in simulated and realistic settings. Furthermore, SpaLP enables zero-shot label transfer on unseen slices, requiring only a small amount of data for pre-training. The reconstructed expression from SpaLP can also enhance the performance of omics foundation model like SCimilarity^30^. We utilized SpaLP to integrate samples from diverse platforms and to dissect multiple complex microenvironments. Taken together, SpaLP provides a powerful tool for ultra-large-scale spatial omics analysis.

Moving forward, several aspects that warrant further exploration include: (1) We plan to further extend SpaLP to incorporate image data for multi-modal integration. Most spatial omics technologies are accompanied by high-resolution imaging, which provides rich histological and morphological information. (2) We also plan to incorporate the integration tasks of unpaired spatial multi-omics data into SpaLP. While a recently published study^77^ has achieved this functionality, it failed to handle ultra-large-scale data due to the limitations of GNN. (3) Omics foundation models are cutting-edge feature representation methods, which improve diverse downstream tasks^72,73,78-80^. However, such models are typically trained on a computing cluster featuring dozens to hundreds of GPUs and huge memory. LP can significantly reduce resource consumption and provide more development opportunities for researchers. Therefore, we plan to develop a multi-modality foundation model through its generalization ability. (4) As a novel and general framework, we plan to extend LP for more large-scale computing fields in the future. Given the generality of the index matrix and the adjacency matrix, LP can naturally substitute GNN.

With the increasing availability of large-scale data, we expect that LP will be widely applied in various fields, advancing the development of computational biology.

## Methods

### Human research ethics approval

This study was approved by the Shanghai Changhai Hospital Medical Ethics Committee (Approval No.: CHEC-2025-045), and the participant voluntarily signed the informed consent form for the institutional sample library. The sharing of the original data was included in and permitted by the initial ethics approval.

### Human kidney cancer CosMx SMI assay

To obtain the human kidney cancer dataset, the Human Universal Cell Characterization RNA 1K-plex panel was used for Nanostring CosMx profiles. Before starting RNA detection cycling, morphology markers for visualization and cell segmentation were added, including pan-cytokeratin (PanCK), CD45, CD298, and DAPI. Over 1037 FOVs (300 μm × 300 μm) were selected for data collection on each slice. The assay employs widefield epifluorescence illumination via a combination of lasers and LEDs, including a 385 nm LED for DAPI, a 488 nm laser for CD298, a 530 nm laser for PanCK, and a 590 nm LED for CD45. Z-stack imaging was performed at each FOV with 7 layers and a step size of 0.5 μm to ensure comprehensive depth coverage.

### Data preprocessing

#### Single sample data preprocessing

For single sample, the genes or proteins expression counts were log-transformed and normalized by library size via the SCANPY^81^ package. We selected top 5,000 highly variable genes (HVGs) for Stereo-seq V1.3^11^ and VisiumHD^10^ platform as input to the SpaLP. For other platforms with gene panels less than 5000, all genes will be used as input.

#### Multi samples data preprocessing

For multi-samples, we retained the common genes or proteins across different samples as input. After selection, each sample was normalized individually. The gene or protein expression counts were log-transformed and normalized by library size using the SCANPY package.

#### Multi-omics data preprocessing

For multi-omics sample, we horizontally concatenated the two omics of each cell into a unified feature as the input.

### Local Pooling framework

Local Pooling (LP) framework is designed to optimize GNN-based methods across various domains involving ultra-large-scale graph-structured data. The framework consists of two distinct modules: (1) Indexing module, (2) Local Pooling module. Compared to traditional GNN-based methods, leveraging its novel architecture, LP demonstrates remarkable computational efficiency requiring substantially fewer computing resources and achieving performance up to several hundred times faster. Moreover, LP eliminates the need for sampling and thus preserves graph integrity on large-scale data, which enables more comprehensive structural learning and more effective neighborhood information aggregation.

### Indexing module

#### Construction of index matrix

For each spatial omics slice *S*, we search the index of the k-nearest neighboring nodes corresponding to each central node. The central nodes and their corresponding neighbor indices are subsequently assembled into an index matrix *I*^(*s*)^. For multiple slices, we concatenate the index matrices of each slice into a disconnected multi-graph index matrix *I*^(1−s)^:

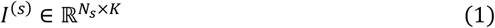

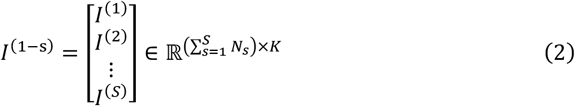

where *N*_*s*_ denotes the number of cells in spatial slice *S*, and *K* denotes the number of nearest neighbors considered for each cell.

Compared with the traditional adjacency matrix, the memory usage of the index matrix increases linearly with the number of cells *N*, while *K* remains constant. In contrast, the adjacency matrix scales quadratically with *N*. As a result, the memory requirement of the adjacency matrix becomes prohibitive when the number of cells reaches hundreds of thousands or even millions, whereas the index matrix can still maintain low memory demands

### Local Pooling module

To encode spatial omics data into a latent space and aggregate neighbor information using the index matrix, we employed an encoder framework built upon the Local Pooling architecture.

#### The input of the local pooling module

For unimodal data 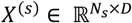 in spatial slice *S*, in which *D* represents the number of proteins or genes in the slice *S* expression matrix. For multiple slices, we concatenate the unimodal data of each slice into feature matrix *X*^(1−s)^:

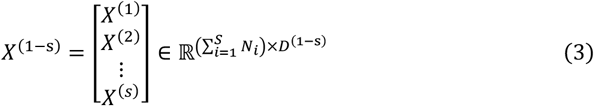

where *N*_*s*_ denotes the number of cells in spatial slice *S, N*_*i*_ denotes the number of cells in spatial slice *i*, and *D*^(1−s)^ denotes genes or proteins that are common to multiple slices. For multimodal data 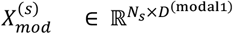 and 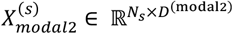 in slice *S*, we define the feature matrix 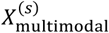 as:

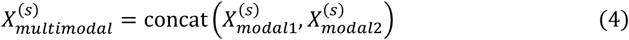

where 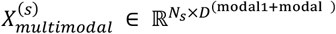 and *D*^(modal2)^ represent the number of features in multimodal, respectively.

#### Local spatial feature

We first encode the feature matrix into the hidden space to obtain the initial latent representation *Z* through a Multilayer Perceptron (MLP). MLP consists of one layer with hidden-dimension size *C*. For cell *i* in the slice *S*, the MLP learns internal cell representations from the full omics feature vector 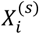 before neighborhood aggregation and reduces the dimensionality:

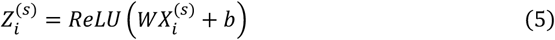

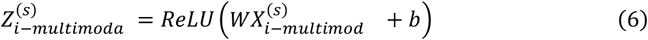

where *W* and *b* are the learnable weight and bias parameters of the linear layer, *ReLU* represents the activation function. The corresponding latent representations of multimodal data is denoted as 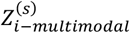.

For cell *i* in the multiple slices, MLP learns internal cell representations from the full omics feature vector before neighborhood aggregation and reduces the dimensionality:

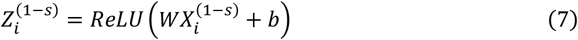

After obtaining the initial latent representation *Z*^(*s*)^ and the index matrix *I*^(*s*)^, the local spatial feature 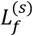 is constructed by gathering the neighboring features of each node from *Z*^(*s*)^ according to the indices in *I*^(*s*)^.

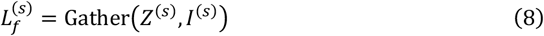

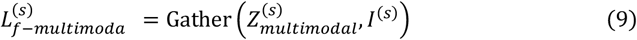

where 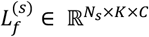 and 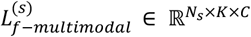. Gather represents extracting neighbor features from *Z*^(*s*)^ by index *I*^(*s*)^.

For multiple slices data, all slices are constructed as a single batch of local spatial features that are not interconnected and the corresponding local spatial feature is denoted as 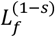:

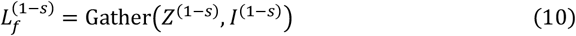

where 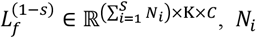 denotes the number of cells in spatial slice *i*.

#### Multi-slice batch integration

When multiple slices are concatenated into a single batch 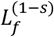 during training, a single Batch Normalization layer is applied to compute channel-wise statistics (mean and variance) across all samples and their local neighborhoods. Shared encoder will align the feature distributions among different slices, resulting in an integrated embedding space where inter-slice batch effects are effectively eliminated.

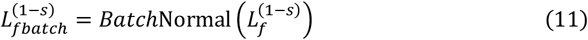

#### Local attention

To effectively aggregate information from the neighbors, we introduce a local attention mechanism. In each feature channel *C*, attention weights are computed across the *K* neighbors to capture their relative importance:

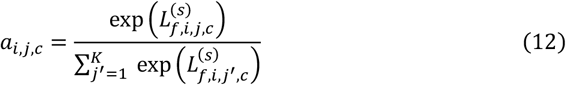

where 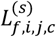 denotes the *C*-th feature of the *j*-th neighbor of the *i*-th center node in the local spatial feature 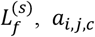 denotes the *C*-th feature of the *j*-th neighbor of the *i*-th center node in the local spatial attention 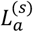.

The local spatial attention 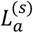 are then used to aggregate local features through a weighted summation:

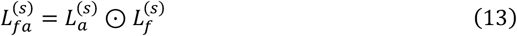

where 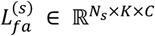 denotes the weighted local feature, 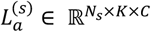 denotes the local attention, and ⊙ denotes Hadamard product. We then aggregate the weighted neighbor features of each central node across all channels to derive its final feature representation 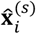. The set of all such representations forms the latent embedding of the entire cell 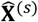.

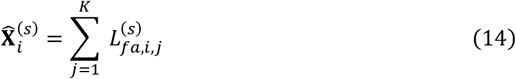

where 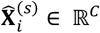 denotes the aggregated feature representation of the *i*-th central node in slice *S*. After model training, the latent representation 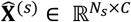 can be used in various downstream analyses, including clustering, visualization and multi-slice batch integration.

### Omics decoder

The decoder reverses the latent embedding of the entire cell back into the expression space. Specifically, for each cell *i* in slice *S*, we use a single-layer linear decoder to process its latent embeddings 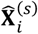 and calculate the reconstruction results 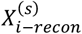 :

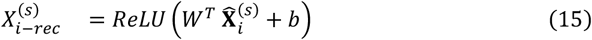

where *W* and *b* are learnable weight matrix and bias vector, *ReLU* activation is conducted to get non-negative reconstruction.

### Model training of Local Pooling

The Local Pooling module is trained to minimize the expression reconstruction loss:

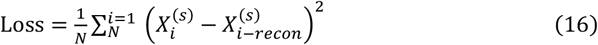

where 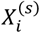 and 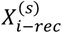 represent the preprocessed original expression features and reconstructed expression feature, respectively.

### Generalization across platform

SpaLP supports direct inference of embeddings on brand-new slices from the same organization without additional training. The newly inferred slice embeddings can be utilized for niche identification and label transfer, where the transferred labels originate from the pre-training dataset. To achieve this, we first trained SpaLP on the pre-training dataset and then froze the parameters. The embeddings of the new data can be directly obtained through SpaLP without additional training. This capability demonstrates the strong generalization power of the LP framework.

#### Label Transfer

SpaLP employs cosine similarity to measure the correspondence between pre-training embeddings 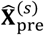 and new data embeddings 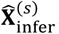. For any cell *i* from the pre-training data, its cosine similarity with cell *j* in the new slice is defined as:

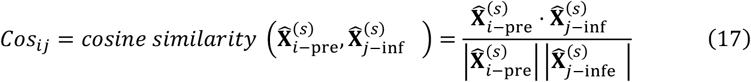

For each cell in the new slice, we assign the label of the nearest cell in the pre-training dataset based on embedding similarity. The cell *i*\* from the pre-training data with the highest cosine similarity is identified as:

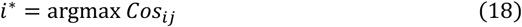

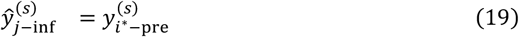

where argmax denotes the operation that returns the cell *i*\* from the pre-training data whose cosine similarity *Cos*_*ij*_ with the new cell *j* is maximal; 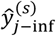 is the predicted label assigned to the new cell, inherited from its most similar cell 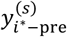 in the pretraining data.

### Experiments

All experiments were performed on a NVIDIA A800-SXM4-80 GB GPU and Intel(R) Xeon(R) Platinum 8462Y+(32 cores, 64 threads) CPU. In all the experiments, we used Adam optimizer with learning rate = 0.001. For 8.4 million cells MERFISH Mouse Brain Atlas, we iteratively input each slice and set the training epoch = 20. For all other experiments, epoch=200. Since all algorithms perform niche identification based on embeddings, the reported running times reflect only the duration required for embedding generation, excluding the time for leiden clustering.

#### Niche identification

Tissue niche identification was performed using the leiden clustering. Specifically, Leiden clustering was applied to the embedding via the scanpy.tl.leiden() function from SCANPY^81^ package. For the ultra-large-scale data of more than 2 million cells, we utilized the GPU version of SCANPY from the rapids-SingleCell and implemented leiden clustering through the function rapids-SingleCell.tl.leiden().

#### Spatial autocorrelation score

We calculated Spatial autocorrelation score with the Squidpy package^82^ to assess the spatial autocorrelation between original and reconstructed gene expression.

### Simulated data

To ensure the fairness of the simulated experiment, we used simulation datasets from HERGAST^28^, which utilized the integrated Human Lung Cell Atlas (HLCA) as the single-cell reference dataset. The HLCA consists of over 2.3 million lung single cells with well-established cell type annotations. Gong et al^28^ manually designed different spatial patterns with complex and refined spatial structures. By extracting different cell types from HLCA to fill these spatial regions and randomly selecting one cell type to be diffusely distributed throughout the entire spatial area to mimic widely present cells. We selected a simulated dataset of 640,000 cells to evaluate the performance of SpaLP and baseline methods on large-scale data.

## Data and detailed methods

Details on the downstream analyses, competing methods and metrics used are available in the Supplementary Information.

## Data availability

All benchmarking data and results are provided with this paper. All h5ad files have been uploaded and are freely available at https://zenodo.org/records/18483604.

## Code availability

The python package source code of SpaLP is available at https://github.com/dbjzs/SpaLP. The jupyter tutorials for reproducing all results in this paper are available at https://spalp.readthedocs.io/en/latest/index.html.

## Acknowledgements

This research was funded by the National Natural Scientific Foundation of China (No. 62571279 (Y.Z.), No. 62303119 (Z.Y.), No. 32470706 (Z.Y.)), the Group Project of Developing Inner Mongolia through Talents (No. 2025TEL25 (Y.Z.)), the Computational Biology Program (No. 25JS2850200 (Z.Y.)) of Science and Technology Commission of Shanghai Municipality (STCSM), and the Central Guidance Fund for Local Science and Technology Development (No. 2024ZY0168 (Y.Z.)).

## Author Contributions Statement

Y.Z. and Z.Y. conceptualized and supervised the project. B.D. designed and developed the method. B.D., Y.L. and L.Y. prepared the figures and tables, contributed to the analysis of data. B.D., Y.L., L.Y., P.H., S.Y., and Y.Z. wrote and revised the manuscript. M.H. and L.W. collected the kidney cancer sample. B.Q. performed Spatial Molecular Imager. All authors have read, revised, and approved the final manuscript.

---

**Extended Data Fig. 1.**
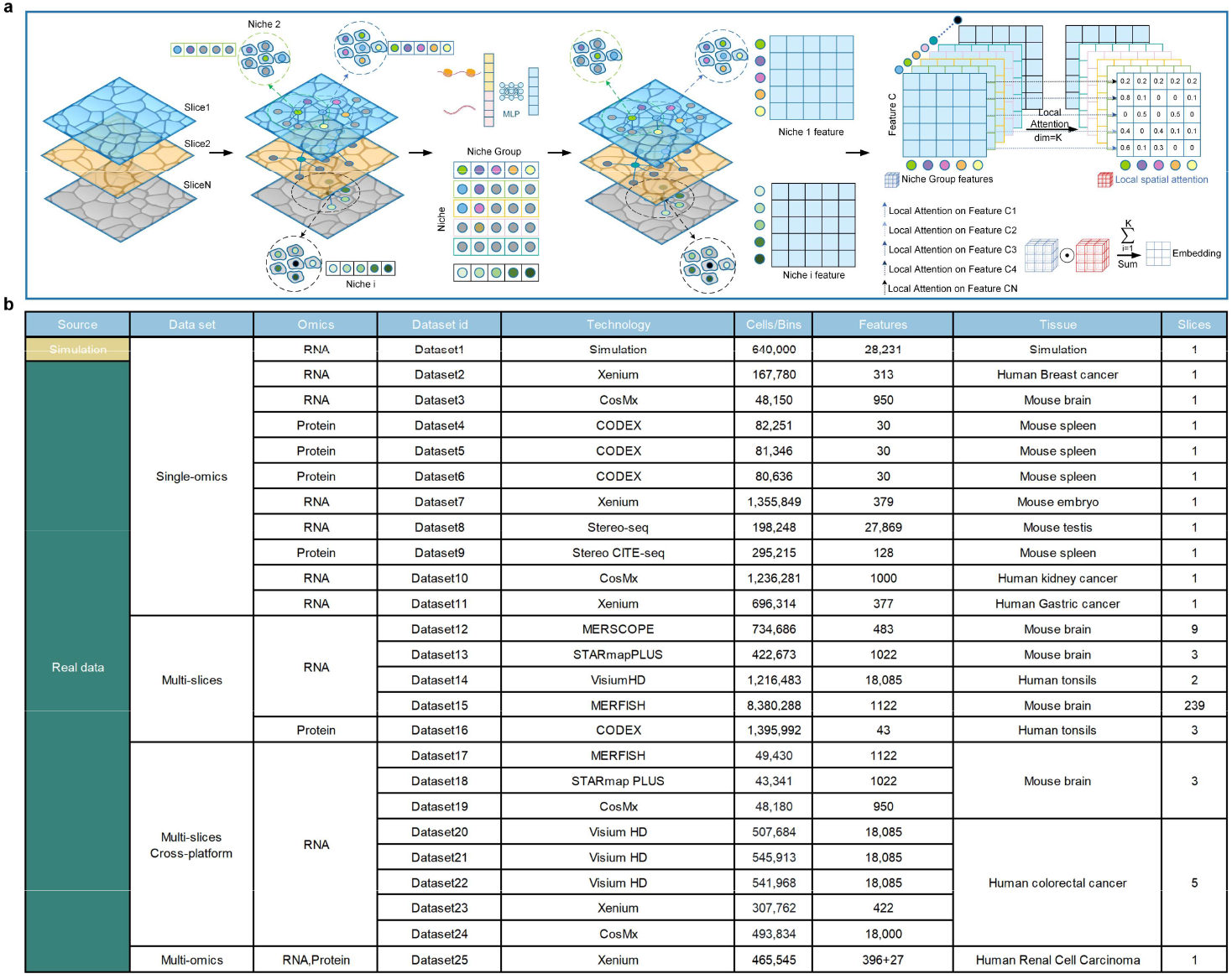
The niche framework of the SpaLP and the datasets involved in the study as well as the statistics. **a**, SpaLP extracts the neighborhood index of each central cell in multiple niches and constructs the niche group. Niche group features are constructed according to the features of each niche, and then the local pooling is used to aggregate the niche features to the central cell. **b**, We comprehensively evaluated SpaLP on over 20 large-scale datasets spanning 9 spatial technology platforms.

**Extended Data Fig. 2.**
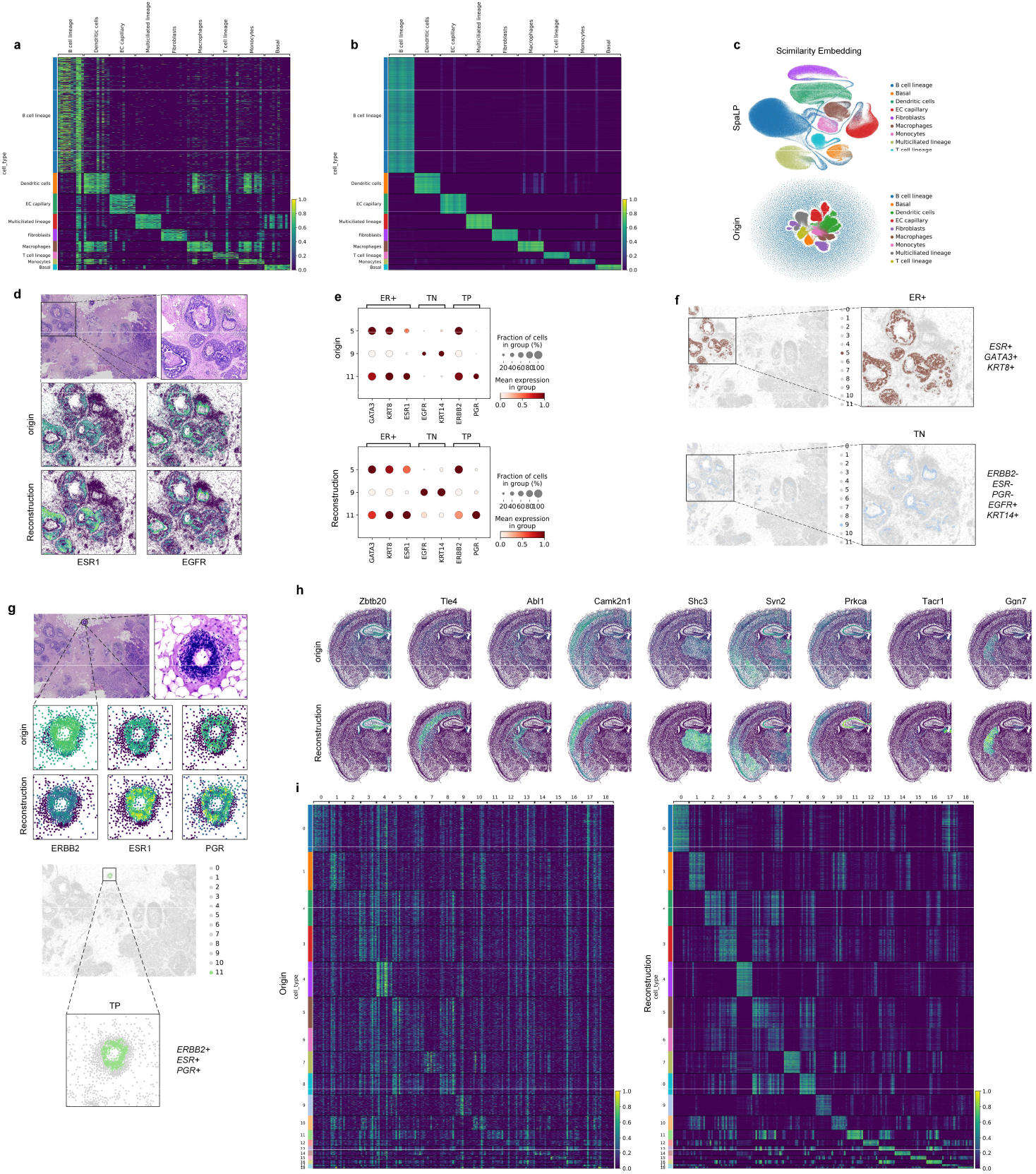
Niches and reconstructed gene expression. **a**, Heat map of original gene expression for each ground truth annotation. **b**, Heat map of reconstructed gene expression for each ground truth annotation. **c**, Embedded-umap visualization from omics foundation model SCimilarity using original and reconstructed gene expression as input. **d**, The original and reconstructed spatial expression of ESR1 and EGFR in DCIS1 region. **e**, Dot plot of the ER+, TN and TP marker genes in SpaLP’s unique niches. **f**, Highlighted visualization of the ER+ and TN in DCIS1 region. **g**, Highlighted visualization of the TP region. **h**, Spatial mapping of original and reconstructed genes expression in CosMx mouse brain. **i**, Heat map of original and reconstructed gene expression in SpaLP’s unique niches.

**Extended Data Fig. 3.**
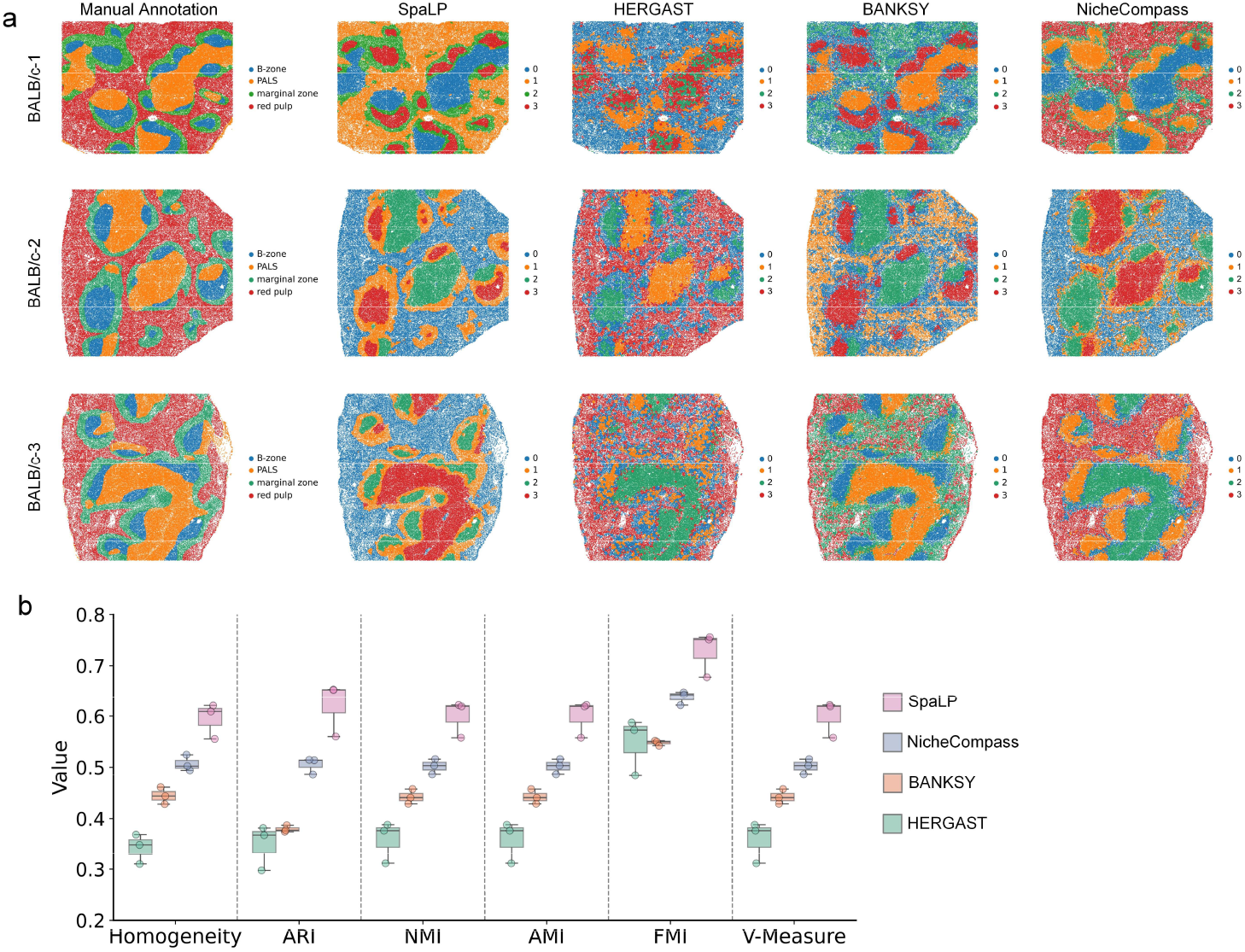
The performance of SpaLP in single-sample niche identification using proteomics. **a**, The manual annotations of three CODEX mouse spleen slices and the corresponding niche identification results from four methods. **b**, Quantitative evaluation of four methods with six supervised metrics.

**Extended Data Fig. 4.**
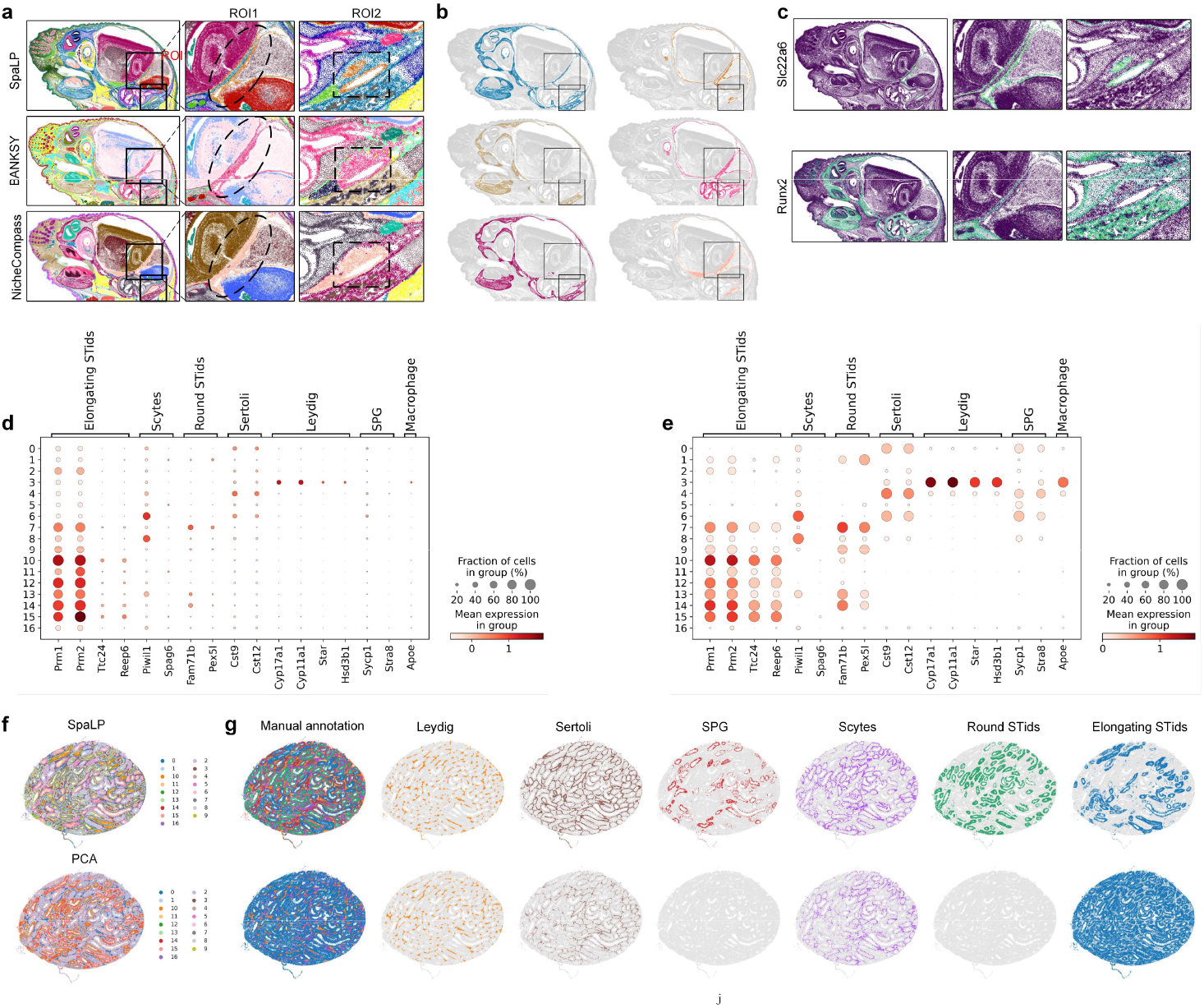
The performance of SpaLP in single-sample niche identification using proteomics. **a**, Zoomed in on the niche of SpaLP and baseline methods in two regions of interest (ROI). **b**, Highlighted visualization of two complementary niches in ROIs. **c**, Zoomed in the spatial mapping of *Slc22a6* and *Runx2* two marker genes. **d**, Dot plot of the original marker genes in SpaLP’s unique niches. **e**, Dot plot of the reconstructed marker genes in SpaLP’s unique niches. **f**, Visualization of the niche identification results for SpaLP and PCA method. **g**, Highlighted visualization of multiple circular gradient niches identification results from SpaLP and PCA.

**Extended Data Fig. 5.**
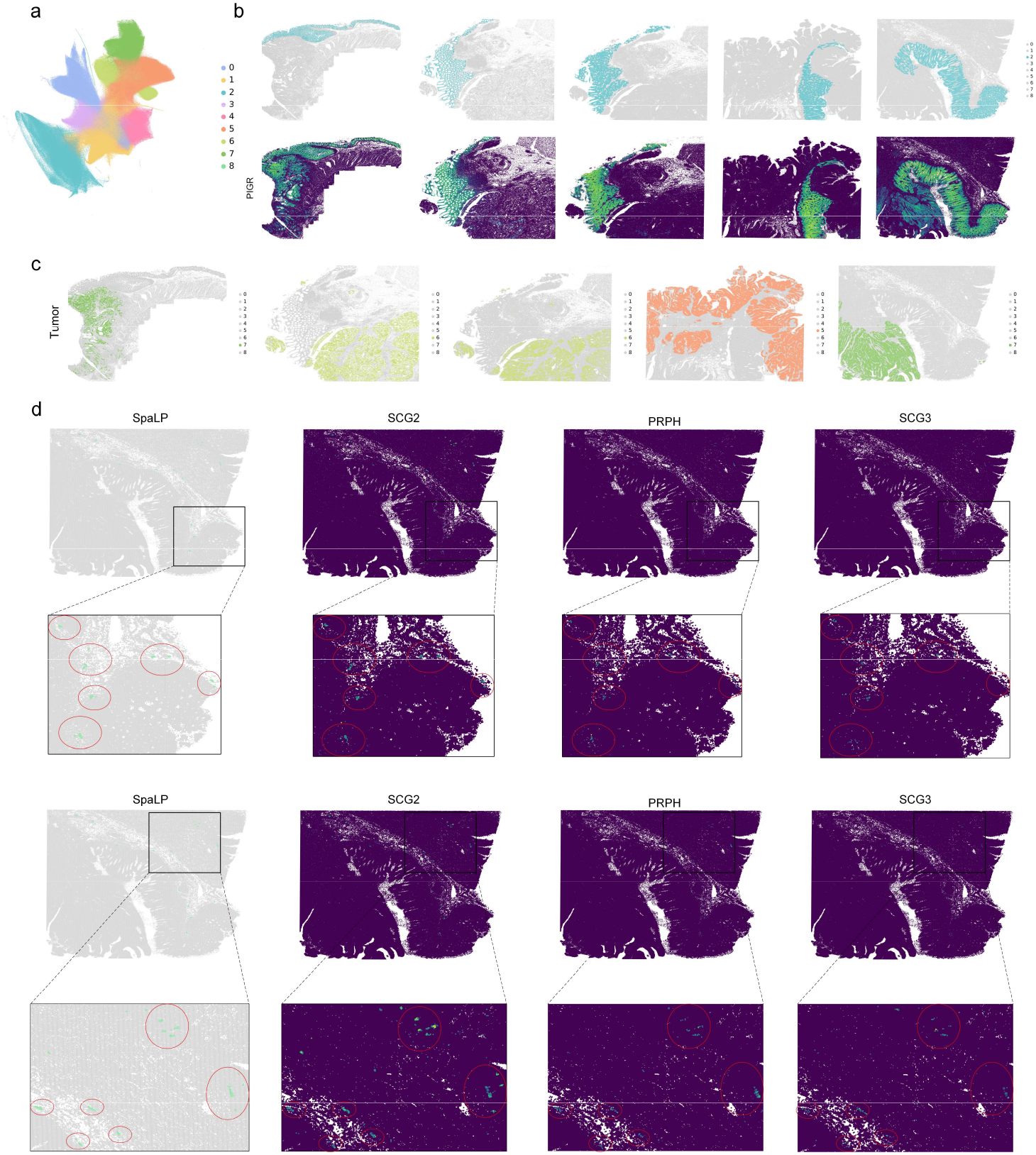
SpaLP effectively integrated conserved niches across slices while preserving slice-specific biological variation and accurately identifying rare niches. **a**, SpaLP’s integrated embedding UMAP and colored by niche identification results. **b**, Highlighted visualization of the goblet cell-rich niche and the corresponding marker genes in five colorectal cancer sections. **c**, Cancer regions across five samples identified by SpaLP. **d**, Highlighted visualization of rare niches and the corresponding marker genes in two regions.

**Extended Data Fig. 6.**
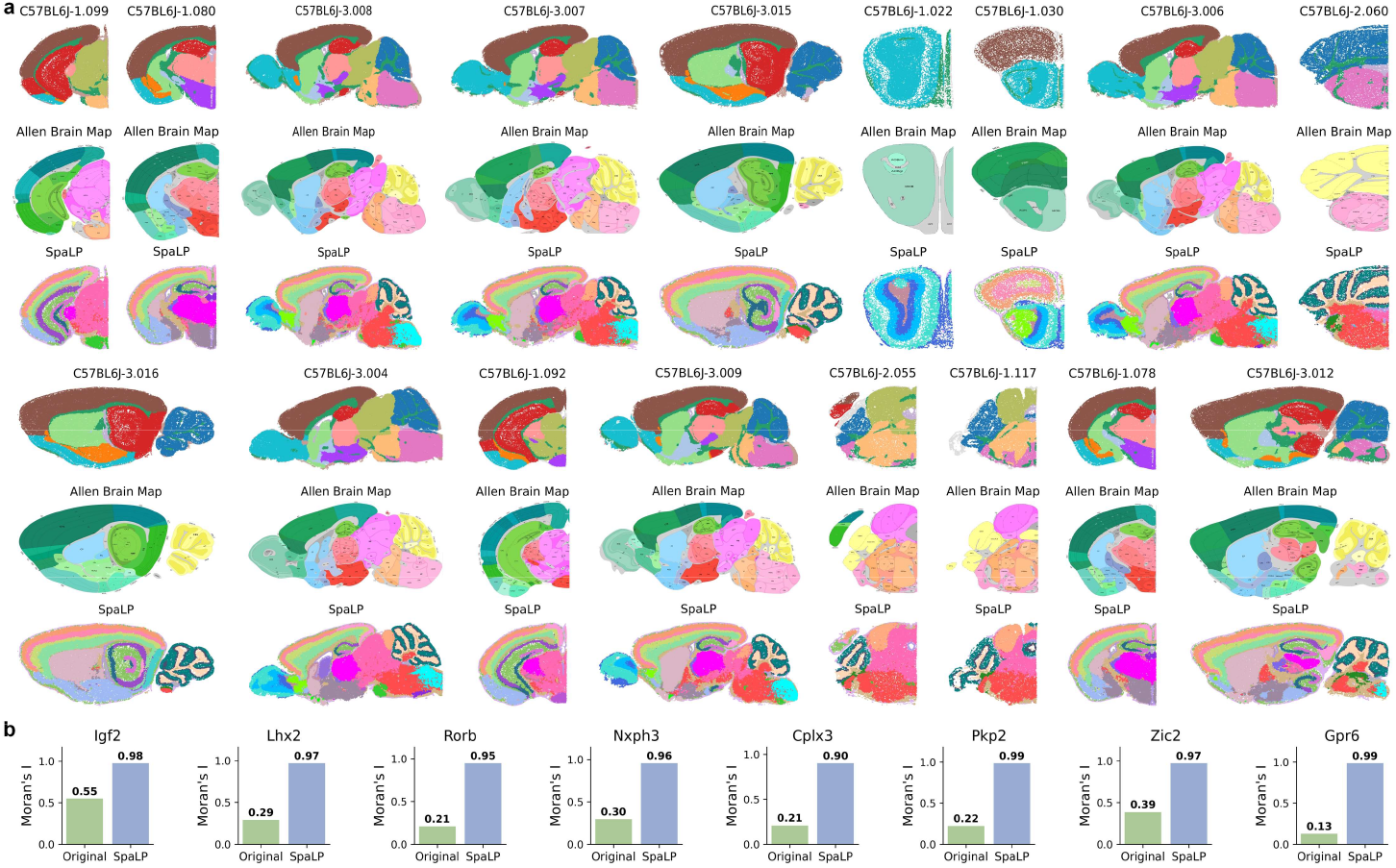
Niches identified in the mouse brain are consistent across slices and correspond to regions from the Allen Brain Map. **a**, Randomly selected tissue slices for all brain region, colored by identified niches and Allen brain annotation. **b**, Bar plots of unsupervised metrics Moran’s I score about the original and reconstructed marker genes.

**Extended Data Fig. 7.**
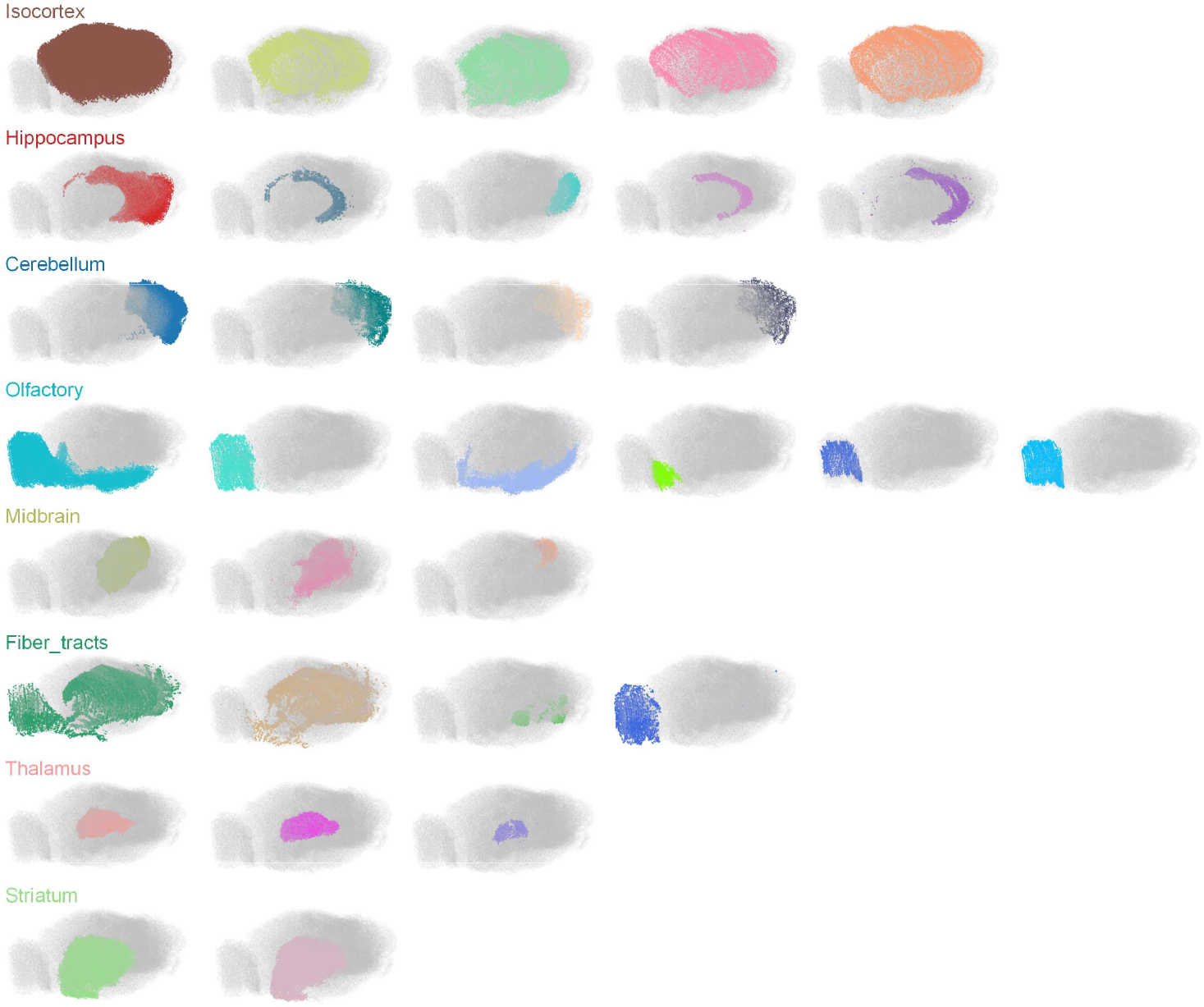
Highlighted visualization of the 3D niche and the corresponding Allen Brain Map.

**Extended Data Fig. 8.**
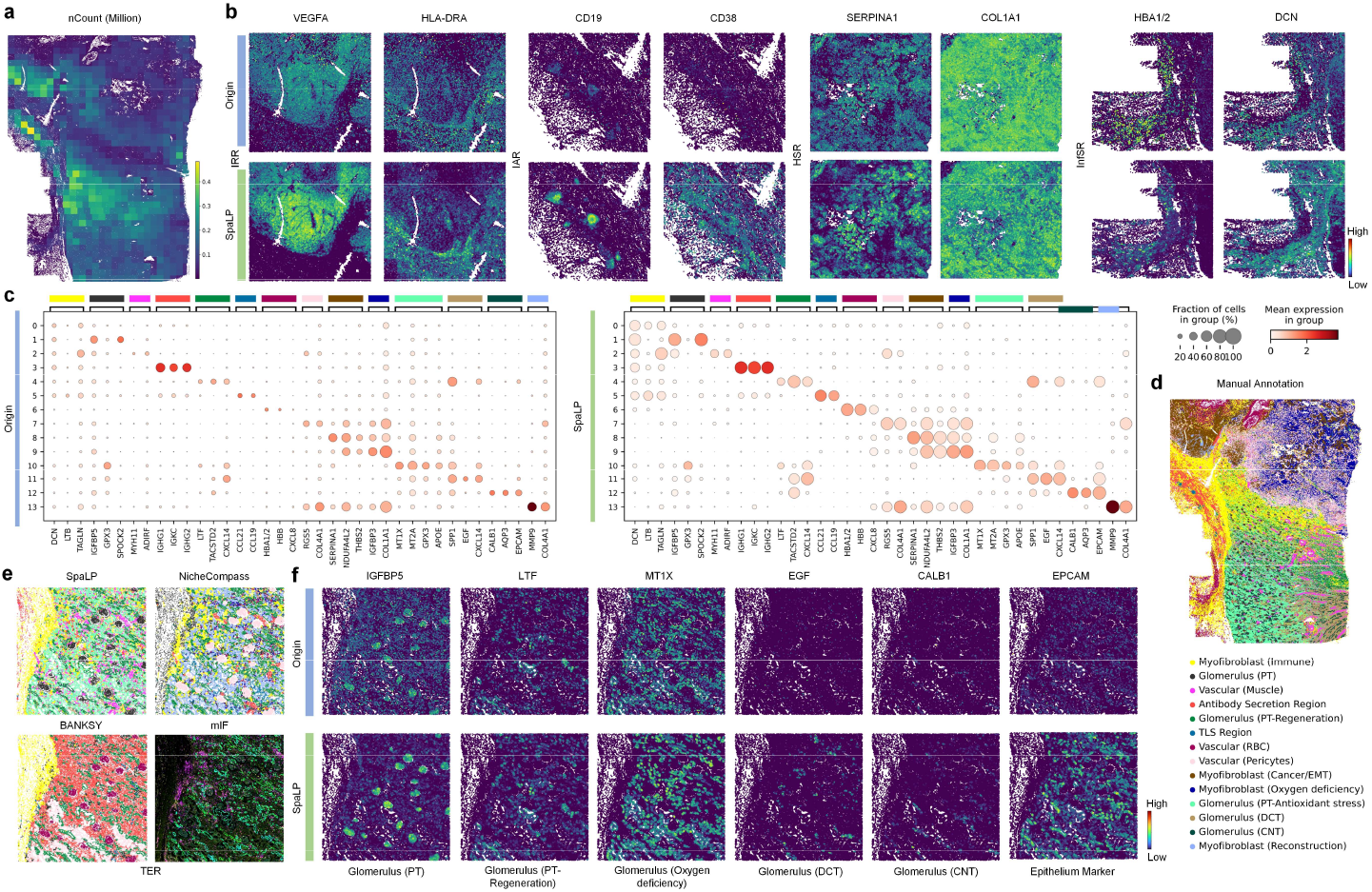
SpaLP maintains its robustness in low-quality kidney cancer sections. **a**, nCount results across 1,037 fields of view (FOVs). **b**, Spatial mapping of original and reconstructed genes in IRR, IAR, HSR and InfSR four regions **c**, Dot plot of the original and reconstructed marker genes in SpaLP’s unique niches. **d**, Manual annotation of the niches identified by SpaLP. **e**, Visualization of the niche identification results from three methods and in TER region and the corresponding marker genes. **f**, Spatial mapping of original and reconstructed expression for multiple marker genes in TER region.

